# Adaptive evolution in virulence effectors of the rice blast fungus *Pyricularia oryzae*

**DOI:** 10.1101/2023.03.16.532886

**Authors:** Marie Le Naour--Vernet, Florian Charriat, Jérôme Gracy, Sandrine Cros-Arteil, Sébastien Ravel, Florian Veillet, Isabelle Meusnier, André Padilla, Thomas Kroj, Stella Cesari, Pierre Gladieux

## Abstract

Plant pathogens secrete proteins called effectors that target host cellular processes to promote disease. Recently, structural genomics has identified several families of fungal effectors that share a similar three-dimensional structure despite remarkably variable amino-acid sequences and surface properties. To explore the selective forces that underlie the sequence variability of structurally-analogous effectors, we focused on MAX effectors, a structural family of effectors that are major determinants of virulence in the rice blast fungus *Pyricularia oryzae*. Using structure-informed gene annotation, we identified 58 to 78 MAX effector genes per genome in a set of 120 isolates representing seven host-associated lineages. The expression of MAX effector genes was primarily restricted to the early biotrophic phase of infection and strongly influenced by the host plant. Pangenome analyses of MAX effectors demonstrated extensive presence/absence polymorphism and identified gene loss events possibly involved in host range adaptation. However, gene knock-in experiments did not reveal a strong effect on virulence phenotypes suggesting that other evolutionary mechanisms are the main drivers of MAX effector losses. MAX effectors displayed high levels of standing variation and high rates of non-synonymous substitutions, pointing to widespread positive selection shaping the molecular diversity of MAX effectors. The combination of these analyses with structural data revealed that positive selection acts mostly on residues located in particular structural elements and at specific positions. By providing a comprehensive catalog of amino acid polymorphism, and by identifying the structural determinants of the sequence diversity, our work will inform future studies aimed at elucidating the function and mode of action of MAX effectors.

**AUTHOR SUMMARY:** Fungal plant pathogens use small secreted proteins, called effectors, to manipulate to their own advantage their host’s physiology and immunity. The evolution of these effectors, whether spontaneously or in response to human actions, can lead to epidemics or the emergence of new diseases. It is therefore crucial to understand the mechanisms underlying this evolution. In this article, we report on the evolution of effectors in one of the prime experimental model systems of plant pathology, *Pyricularia oryzae*, the fungus causing blast diseases in rice, wheat, and other cereals or grasses. We further characterize in this fungus a particular class of effectors, called MAX effectors, using structural models based on experimental protein structures of effectors. We show that this class of effector is produced by the pathogen during the early stages of infection, when plant cells are still alive. By comparing the gene content of isolates infecting different plant species, we show that the MAX effector arsenal is highly variable from one isolate to another. Finally, using the inferential framework of population genetics, we demonstrate that MAX effectors exhibit very high genetic variability and that this results from the action of natural selection.

## INTRODUCTION

Plant pathogens secrete effector proteins to manipulate the physiology and metabolism of their host and to suppress its immunity. Consequently, effectors are expected to engage in coevolutionary interactions with plant defense molecules. The proximate mechanisms of effector-driven adaptation are relatively well-characterized: plant pathogens adapt to new hosts through changes in effector repertoires and effector sequences [1, 2]. However, the ultimate (eco-evolutionary) mechanisms underlying effector diversification have remained elusive. The concept of coevolution posits that adaptation in one partner drives counter-adaptations in the coevolving partner [3-5]. Under the co-evolutionary arms race model, variation for disease resistance and pathogen virulence is transient, resulting in a turnover of sequence variation through repeated episodes of strong directional selection [6]. In agricultural systems, because pathogens tend to be ahead of their hosts in the arms race owing to their larger populations and shorter generation times, the co-evolutionary arms race tends to result in so-called boom and bust cycles [7]. Under the alternative, ‘trench warfare’ hypothesis, advances and retreats of resistance or virulence genes frequencies maintain variation as dynamic polymorphisms [8, 9]. The maintenance of genetic polymorphisms is called ‘balancing selection’, a process by which different alleles or haplotypes are favored in different places (via population subdivision) and/or different times (via frequency-dependent negative selection). While there is a growing body of data demonstrating the nature and prevalence of the selective pressures that shape the diversity of immune systems in plants [6, 10-12]. we still lack a clear picture of the co-evolutionary mechanisms underlying the molecular evolution of virulence factors in their interacting antagonists [13].

Effectors from plant pathogenic fungi are typically cysteine-rich secreted proteins smaller than 200 amino acids with an infection-specific expression pattern. Effectors are numerous in fungal genomes (several hundred to more than a thousand per genome), and rarely show homologies with known proteins or domains. They are also highly variable in sequence and do not form large families of sequence homologs. Based on similarity analyses, fungal effectors can form small groups of paralogs (typically with less than five members), but they are most often singletons. This apparent lack of larger effector families has hindered attempts to probe into the evolutionary factors underlying their diversification. In addition, the high diversity of fungal effectors has hampered functional analyses due to the lack of good criteria for prioritizing them and our inability to predict their physiological role. Consequently, the virulence function and evolutionary history of most fungal effectors remain unknown.

Recently, the resolution of the three-dimensional (3D) structure of fungal effectors combined with Hidden Markov Model (HMM) pattern searches and structure modeling revealed that fungal effector repertoires are, despite their hyper-variability, dominated by a limited number of families gathering highly sequence-diverse proteins with shared structures and, presumably, common ancestry [14-17]. One such structurally-conserved but sequence-diverse fungal effector family is the MAX *{Magnaporthe* Avrs and ToxB-like) effector family. MAX effectors are specific to ascomycete fungi and show massive expansion in *Pyricularia oryzae* (synonym: *Magnaporthe oryzae)* [17]. the fungus causing rice blast disease, one of the most damaging diseases of rice [18, 19]. MAX effectors are characterized by a conserved structure composed of six β-strands organized into two antiparallel β-sheets that are stabilized in most cases by one or two disulfide bridges. The amino acid sequence of MAX effectors is very diverse and they generally have less than 15% identity, which suggests that they are a family of analogous, rather than homologous, effector proteins. In other words, they share a similar three-dimensional structure, but no conclusions can be drawn with respect to shared ancestry. MAX effectors are massively expressed during the biotrophic phase of infection, suggesting an important role in disease development and fungal virulence [17]. Remarkably, about 50% of the known avirulence (AVR) effectors of *P. oryzae* belong to the MAX family, indicating that these effectors are closely monitored by the host plant immune system [17].

*Pyricularia oryzae* is a multi-host, poly-specialist pathogen that infects more than 50 monocotyledonous plants, including major cereal crops such as rice, maize, wheat, or barley [20-23]. *Pyricularia oryzae* has repeatedly emerged on new hosts [21, 24]. in new geographical areas [25, 26]. and phylogenomic analyses have revealed that it can be subdivided into several genetic lineages, each preferentially associated with a specific or restricted set of host plant genera [27]. In *P. oryzae*, effectors can play a major role in host-shifts or host-range expansions [28-30]. For example, loss of function of the PWT3 effector in *Lolium*-infecting strains contributed to gain of virulence on wheat [29]. Similarly, loss of the MAX effector AVR1-CO39 is thought to have contributed to the emergence of rice blast from foxtail-millet infecting isolates [20, 31]. This indicates that MAX effectors may be important determinants of host specificity in *P. oryzae*.

In this study, we characterized the genetic diversity of MAX effectors in *P. oryzae* and within its different host-specific lineages. We explored the evolutionary drivers of the diversification of MAX effectors and tested whether MAX effectors represent important determinants of *P. oryzae* host specificity. To this aim, we assembled and annotated 120 high-quality *P. oryzae* genomes from isolates representing seven main host-specific lineages. We mined these genomes for putative effectors and used hidden Markov models based on fold-informed protein alignments to annotate putative MAX effectors. We identified 58 to 78 putative MAX effector genes per individual genome distributed in 80 different groups of MAX homologs. We showed that the expression of *MAX* effector genes is largely restricted to the early biotrophic phase of infection and strongly influenced by the host plant. Our evolutionary analyses showed that MAX effectors harbor more standing genetic variation than other secreted proteins and non-effector genes, and high rates of non-synonymous substitutions, pointing to positive selection as a potent evolutionary force shaping their sequence diversity. Pangenome analyses of MAX effectors demonstrated extensive presence/absence polymorphism and identified several candidate gene loss events possibly involved in host range adaptation. Our work demonstrates that MAX effectors represent a highly dynamic compartment of the genome of *P. oryzae*, likely reflecting intense co-evolutionary interactions with host molecules.

## RESULTS

### Genome assembly and prediction of MAX effector genes

We used a collection of genome assemblies that included 120 haploid isolates of *Pyricularia oryzae* fungi from 14 different host genera: *Oryza* (n=52), *Triticum* (n=21), *Lolium* (n=12), *Setaria* (n=8), *Eleusine* (n=8), *Echinochloa* (n=4), *Zea* (n=4), *Bromus* (n=2), *Brachiaria* (n=2), *Festuca* (n=2), *Stenotaphrum* (n=2), *Eragrostis* (n=1), *Hordeum* (n=1), and *Avena* (n=1) (S1 Table). Assembly size ranged from 37Mb to 43.2Mb, with an average size of 40.2 Mb (standard deviation [s.d.]: 1.9Mb). L50 ranged from five to 411 contigs (mean: 97.1; s.d.: 83.2) and N50 from 28Kb to 4.0Mb (mean: 238.6Kb; s.d.: 43.8Kb; S1 Table). Gene prediction based on protein sequences from reference 70-15 and RNAseq data identified 11,520 to 12,055 genes per isolate (mean: 11,763.2; s.d.: 103.7). The completeness of assemblies, as estimated using BUSCO [32]. ranged between 93.4 and 97.0% (mean: 96.4%; s.d.: 0.6%; S1 Table).

MAX effectors were identified among predicted secreted proteins using a combination of similarity searches [33, 34] and structure-guided alignments [35] as summarized in Figure 1. To assess variation in the MAX effector content of *P. oryzae*, we constructed groups of homologous genes (i.e., “orthogroups” or OG) using the clustering algorithm implemented in ORTHOFINDER [36]. A given orthogroup was classified as secreted proteins or MAX effectors if 10% of sequences in the group were identified as such by functional annotation. Sequences were grouped in 14,767 orthogroups, of which 80 were classified as encoding MAX effectors and 3,283 as encoding other types of secreted proteins (Figure 1). The number of MAX orthogroups per isolate ranged from 58 to 73 (average: 65.8; s.d.: 2.8), representing between 58 to 78 *MAX* genes per isolate (average: 68.4; s.d.: 3.6). The 80 orthogroups of MAX effectors were further split into 94 groups of orthologs, by identifying paralogs using gene genealogies inferred with RAxML v8 [37] (S2 Table). Comparison of these 94 groups of orthologs with MAX effectors predicted by previous studies [15-17, 38] revealed that 19 were not predicted by any other study, while 75 were predicted by at least one other study (including nine predicted by all studies). Twenty-five MAX effectors predicted by other analyses, were not identified by our prediction pipeline and were, therefore, not considered in our study (S2 Table).

**Figure 1.**
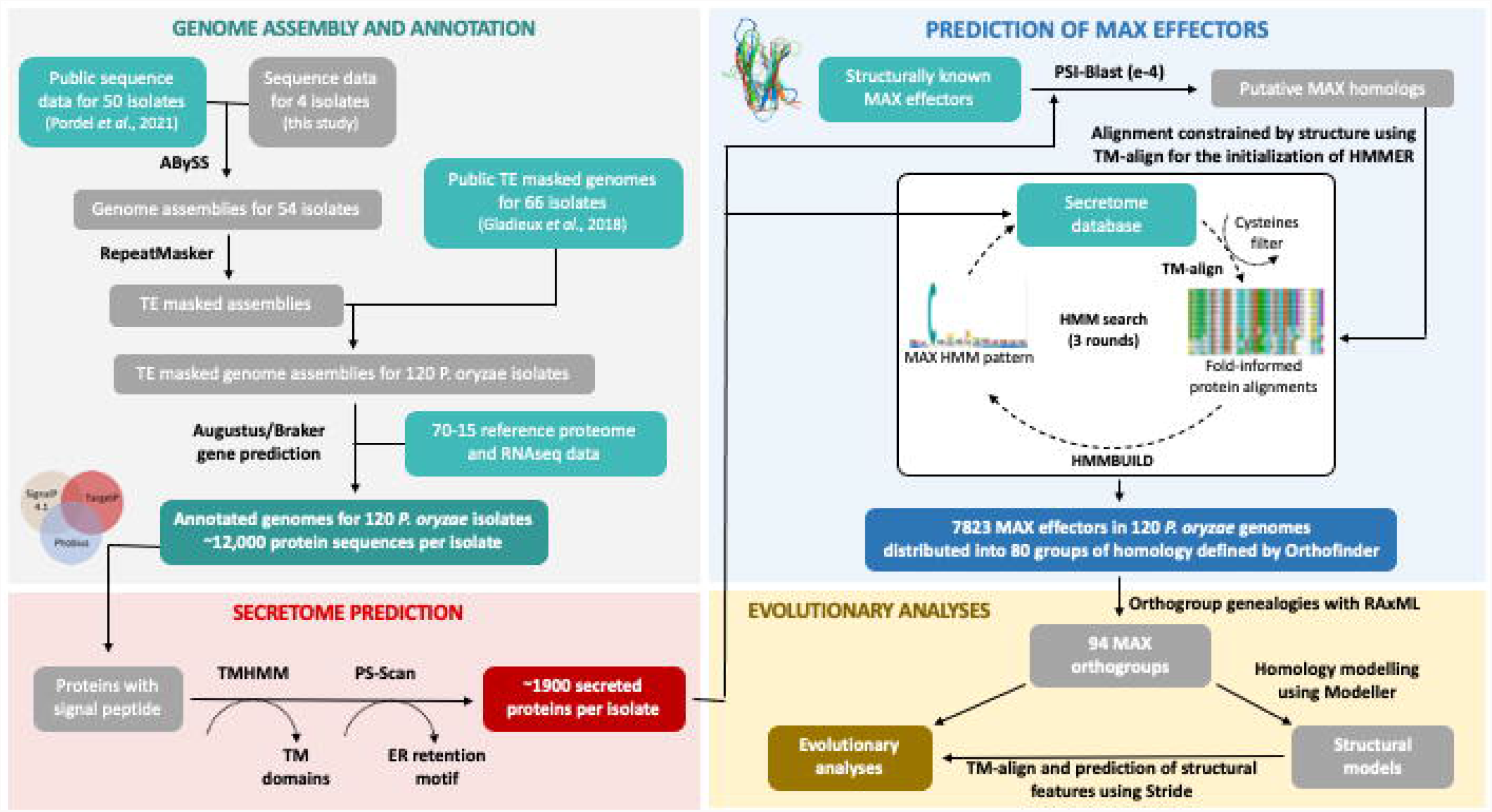
Schematic representation of the main steps of the bioinformatic pipeline used to predict genes in 120 genomes of *Pyricularia oryzae*, to identify genes encoding MAX effectors, and to measure their genetic variability. Gladieux et al. 2018 [27]; Pordel et al. 2021 [21]. Genome assembly and annotation: using RNAseq and reference proteins, we predicted genes with AUGUSTUS 3.4.0 [56] and Braker 1 [70] in 54 genomes that were assembled in this study with ABYSS 1.9.0 [69]. as well as in 66 assemblies that were published earlier; repeated sequences were masked using REPEATMASKER 4.1.0 (http://www.repeatmasker.org/). Secretóme prediction: we identify signal peptides in the predicted genes by running Signal P 4.1 [74]. Target P 1.1 [75]. and Phobius 1.01 [62]. removing genes encoding proteins with a predicted transmembrane domain (identified using Tmhmm [76]), as well as endoplasmic reticulum proteins (identified with PS-SCAN from expasy.org). Prediction of MAX effectors: MAX effectors were identified using PSI-BLAST [33] to search for homologs of known MAX effectors (AVR1-CO39, AVR-Pia, AvrPiz-t, AVR-PikD, and ToxB) in the predicted secretóme, followed by structure-guided alignment of PSI-BLAST hits using TM-ALIGN [35]. and three iterative rounds of HMMER [34] searches based on Hmmbuild models built on TM-ALIGN alignments of significant hits. Only proteins with two expected conserved cysteines less than 33-48 amino acids apart were retained [17]. Evolutionary analyses: the 11 orthogroups that included paralogous copies of MAX effectors were split into sets of orthologous sequences using genealogies inferred using RAxML v8 [37]. yielding a total of 94 single-copy MAX orthologs for evolutionary analyses; protein structure models were inferred using homology modeling in MODELLER [79]. and structural features were identified with STRIDE [91].

The number of MAX effectors per isolate was primarily determined by the host of origin. We did not observe any significant relationship between the number of MAX effectors and assembly properties (assembly size and N50), unlike the number of secreted proteins and total number of genes (S1 Figure). Analysis of variance revealed that the host of origin had a significant impact on the number of MAX effectors (F_13,95_=10.33; *p*=3e^-8^), while the origin of genomic data did not have an effect (i.e., the study in which genomic data were initially described; F_11,95_=1.07; *p*=0.39; S1 Figure). These analyses indicate that the observed variation in the size of the MAX effector repertoire is primarily biological in origin, not technical.

#### MAX effectors are massively deployed during rice infection

To determine whether these putative MAX effectors are deployed by *P. oryzae* during plant infection, we analyzed the expression patterns of the 67 *MAX* genes predicted in the genome of the reference isolate Guy11 by qRT-PCR (Figure 2A). Using RNA samples from Guy11 mycelium grown on artificial media, we found that 94% of the *MAX* genes (63 genes) were not, or very weakly expressed during axenic culture, and only four (i.e., *MAX24, MAX29, MAX59*, and *MAX66)* showed weak, medium or strong constitutive expression (Figure 2 A). *MAX* genes were, therefore, predominantly repressed in the mycelium of *P. oryzae*.

**Figure 2.**
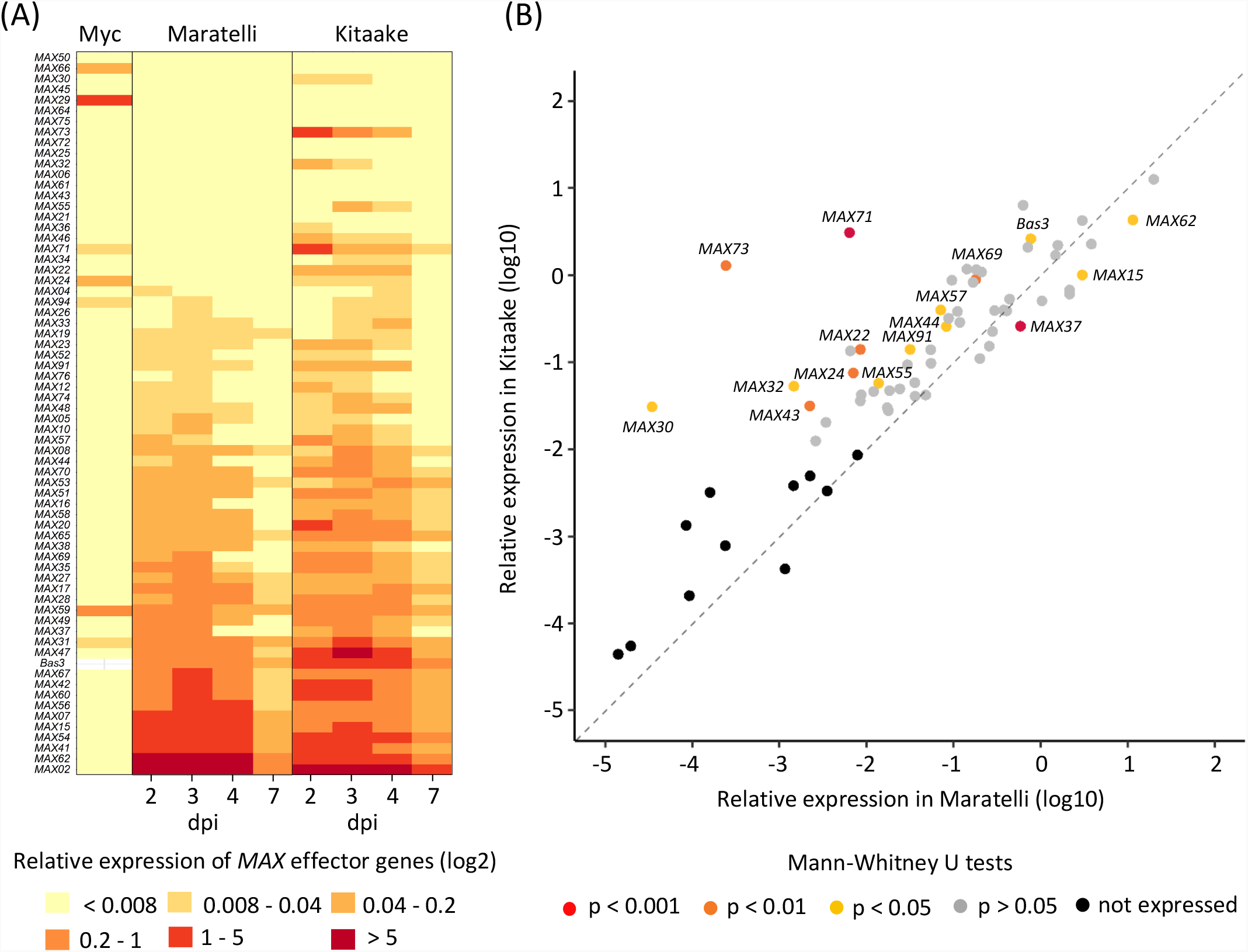
The expression of *MAX* genes is biotrophy-specific and is influenced by the host plant. (A) Transcript levels of *MAX* genes and the biotrophy marker gene *Bas3* were determined by qRT-PCR in the mycelium (Myc) of the *P. oryzae* isolate Guy11 grown for 5 days in liquid culture, and in infected leaves of the rice cultivars Maratelli and Kitaake at 2, 3, 4, and 7 days post inoculation (dpi) with Guy11. Relative expression levels were calculated using the constitutively expressed *MoEFlα (Elongation Factor 1α)* gene as a reference. The heatmap shows the median relative expression value for each gene (in log2 scale), calculated from 6 independent biological samples for the Myc condition, and 3 independent inoculation experiments (each with 5 independent leaf samples per time point) for each rice cultivar. Effectors were ranked from top to bottom by increasing relative expression values in Maratelli. Relative expression values were assigned to six categories: not expressed (<0.008), very weakly (0.008-0.04), weakly (0.04-0.2), moderately (0.2-1), strongly (1-5) and very strongly expressed (>5). (B) Scatter plot comparing the relative expression levels of *MAX* genes in Guy11-infected Maratelli and Kitaake cultivars. Each point shows the maximum median relative expression value (in log10 scale) calculated in the infection kinetics described in (A). Difference in effector relative expression levels between the two conditions was assessed by Mann-Whitney U tests and dots were colored according to significance results: grey (*p*>0.05), yellow (*p*<0.05), orange (*p*<0.001), red (*p*<0.0001), black (effectors not expressed in both conditions).

We analyzed the expression of the *MAX* genes 2, 3, 4, and 7 days following spray inoculation of Guy11 on the rice cultivar Maratelli, which is highly susceptible to *P. oryzae*. We found that 67% of the *MAX* genes (45 genes) were expressed (Figure 2A). Among them, three were also expressed in the mycelium (i.e., *MAX31, MAX59*, and *MAX94). MAX31* was over-expressed under infection conditions, whereas the other two showed similar expression levels *in vitro* and during infection. 64% of the *MAX* genes (42 genes) showed an infection-specific expression profile with relative expression levels ranging from very low (0.008-0.04) to very high (>5). Like the *Bas3* gene, encoding a *P. oryzae* effector specifically induced during the biotrophic phase of infection, all *MAX* genes showed maximal expression between the second and fourth day post-inoculation (Figure 2A; S2 Figure).

To test whether the genotype of the host plant could influence the expression of *MAX* genes, we analyzed their expression patterns upon infection of the rice cultivar Kitaake, which has a higher basal resistance to *P. oryzae* than Maratelli. We computed the median relative expression across three independent experiments with five biological replicates each (Figure 2A; Figure 2B). During Kitaake infection, 78% of the *MAX* genes (52 genes) were upregulated compared to the *in vitro* condition, while 64% (43 genes) were induced upon infection of Maratelli (Figure 2A). Some *MAX* genes not expressed in Maratelli were induced in Kitaake (e.g., *MAX24, MAX30, MAX32, MAX43, MAX71*, and *MAX73*) (Figure 2B, S3 Figure). Others were significantly upregulated in Kitaake compared to Maratelli (i.e., *MAX22, MAX44, MAX55, MAX57, MAX69*, and *MAX91*). However, a few genes, such as *MAX15, MAX37* and *MAX62*, among the most strongly expressed effectors in Maratelli, showed weaker levels of expression in Kitaake. These results show that Guy11 deploys a wider diversity of MAX effectors during the infection of Kitaake compared to that of Maratelli, and that MAX effectors are subject to host-dependent expression polymorphism.

Taken together, our data indicate that during the biotrophic phase of rice infection, *P. oryzae* actively expresses a significant portion of its MAX effector repertoire in a host-dependent manner, which suggests that these effectors have an important function in fungal virulence.

### The MAX effector repertoire is highly variable

To investigate the genetic diversity of MAX effectors in *P. oryzae*, we analyzed their nucleotide diversity per base pair (π), their ratio of non-synonymous to synonymous nucleotide diversity (*π*_*N*_/*π*_*S*_, page 226 in ref. [39]), and their presence-absence polymorphism. Compared to other secreted proteins or other genes, MAX effector orthogroups had higher *π*, and *π*_*N*_*/π*_*S*_ values, and lower presence frequency (S4 Figure). Orthogroups including known avirulence genes like *AVR1-C039, AvrPiz-t* and *AVR-Pik* featured among the most diverse orthogroups of MAX effectors (S1 Data).

We categorized genes in the pangenome according to their presence frequencies [36]. with core genes present in all isolates, softcore genes present in >99% isolates, shell genes present in 1-99% isolates and cloud genes present in <1% isolates. The majority of MAX effector genes were classified as shell (64/80 [80%] orthogroups), while the majority of other secreted proteins or other genes were classified as core or softcore (1650/3283 [50.2%] and 6714/11404 [58.9%] orthogroups, respectively) (Figure 3A). Only a minority of genes were present in multiple copies (MAX: 15/80 [18.8%]; other effectors: 746/3283 [22.7%]; other genes: 1439/11404 [12.6%]; Figure 3A). Assessment of the openness of the pan-genome by iteratively subsampling isolates revealed a closed pangenome with a limited number of pan and core genes for MAX effectors, other secreted proteins and the remainder of the gene space (Figure 3B). Nucleotide diversity differed significantly between categories of the pangenome for non-MAX effectors (Kruskal-Wallis test: H=181.17, d.f.=2, *p<0*.001) and other genes (Kruskal-Wallis test: H=225.25, d.f.=2, *p<*0.001), but not for MAX effectors (Kruskal-Wallis test: H=2.50, d.f.=2, *p*>0.05). For non-MAX effectors and other genes, nucleotide diversity π was significantly higher in the shell genes than in softcore genes and core genes (Post-hoc Mann-Whitney U-tests, p<0.001; Figure 3C). The frequency of MAX orthogroups was positively and significantly correlated with the frequency of neighboring orthogroups at the species-wide level and at the level of Oryza and Setaria lineages, which indicates that non-core MAX effectors tend to be located in regions with presence-absence variation (S5 Figure).

**Figure 3.**
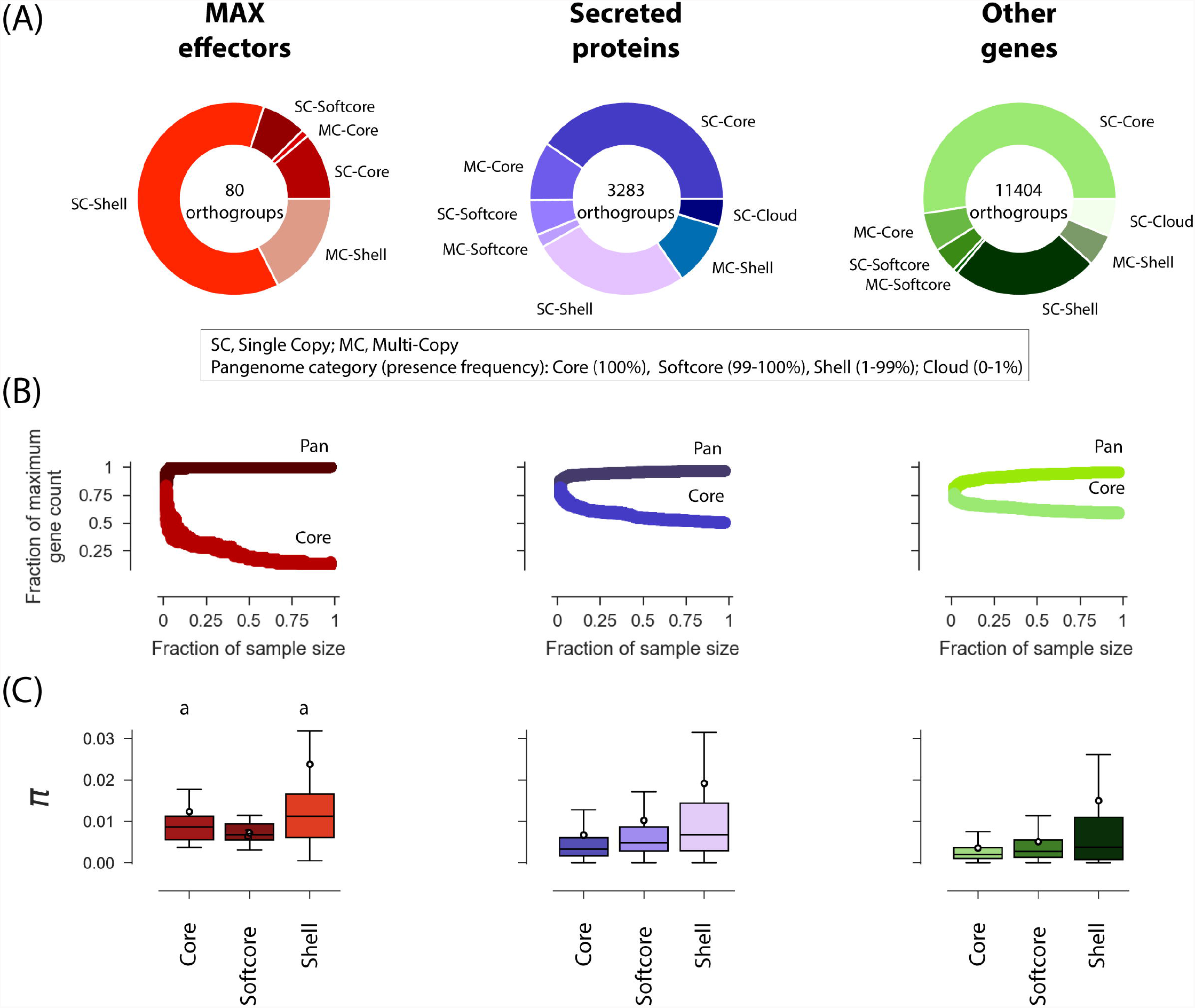
The pan-genome of *P. oryzae*. (A) Composition of the pangenome of MAX effectors, other secreted proteins, and other genes. (B) Rarefaction analysis of the size of pan- and core-genomes. For k ranging from two to the sample size minus one, pan- and core-genome sizes were computed for 1000 random combinations of k isolates. Subsample size is represented as a fraction of the sample size (n=120), and pan- and core-genome sizes as a fraction of maximum gene counts (reported at the center of donut plots in panel A), “core” genes are present in all isolates of a pseudo-sample of size k; “pan” qualifies genes that are not “core”. (C) Nucleotide diversity per base pair *(π)* in core, softcore, and shell genes. A number of data points were cropped from the nucleotide diversity plot for visually optimal presentation but included in statistical tests. In box plots, the black circle is the mean, the black line is the median. Cloud genes were not included in the nucleotide diversity plot because it was not computable due to the small sample size or lack of sequence after filtering for missing data. Shared superscripts indicate non-significant differences (Post-hoc Mann-Whitney U-tests).

Together, these analyses show that the MAX effector repertoire is highly plastic compared to other gene categories, both in terms of the presence/absence of orthogroups and the sequence variability within orthogroups.

### MAX effector variability is structured by host plant

To investigate signatures of positive selection in the genome of *P. oryzae*, and identify candidate loci involved in host specificity, we first identified the divergent lineages represented in our dataset. We inferred population structure from 6780 SNPs in single-copy core orthologs to minimize the potential impact of natural selection on our findings. We used complementary approaches that make no assumption about random mating or linkage equilibrium. Both clustering analyses with the SNMF software [40] (Figure 4A) and neighbor-net phylogenetic networks [41] (Figure 4B) revealed consistent patterns that split genetic variation primarily by host of origin. Although the lowest cross-entropy value was observed at K=11 in the SNMF analysis, we chose to represent K=8 because the cross-entropy was only slightly higher and K=8 did not split the *Triticum-* and *Eleusine-associated* clusters (S6 Figure). Lineage-level analyses were limited to the lineages with the largest sample size, associated with rice ( *Oryza)*, foxtail millet *(Setaria)*, wheat *(Triticum)*, ryegrass *(Lolium)*, and goosegrass *(Eleusine)*.

**Figure 4.**
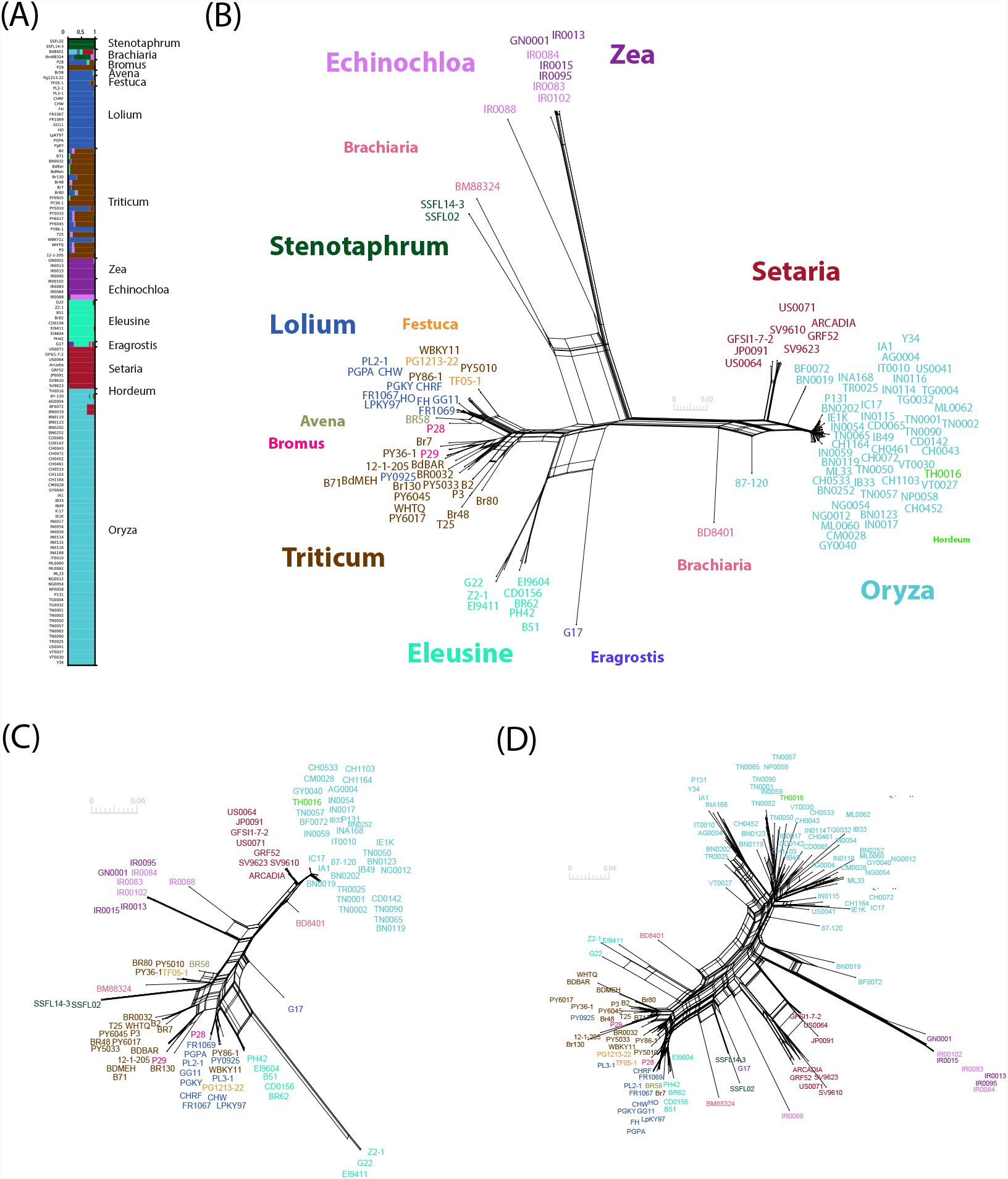
Population subdivision in 120 isolates of *Pyricularia oryzae*. Population subdivision was inferred from (A-B) 6780 polymorphisms at four-fold degenerate synonymous sites identified in coding sequences of single-copy core orthologs (one polymorphism without missing data randomly chosen per ortholog), (C) 130 SNPs without missing data identified in coding sequences of single-copy core MAX effectors, and (D) a table of presence/absence data (coded as 0 and 1) for all 80 MAX effector orthogroups. (A) Genetic ancestry proportions for individual isolates in K=8 ancestral populations, as estimated using the SNMF clustering algorithm [40]; each isolate is depicted as a horizontal bar divided into K segments representing the proportion of ancestry in K=8 inferred ancestral populations; host genus of origin is indicated on the right side. (B-D) Neighbor-net phylogenetic networks estimated with SplitsTree [41]. with isolate names colored according to their host of origin.

Population subdivision inferred from MAX effectors using either 130 SNPs without missing data in single-copy core MAX effectors (Figure 4C) or presence/absence variation of all 80 MAX effector orthogroups (Figure 4D) revealed essentially the same groups as the analysis of the single-copy core orthologs. This indicates that genome-wide nucleotide variation, variation in MAX effector content, and nucleotide variation at MAX effectors reflected similar genealogical processes. The *Oryza* and *Setaria* lineages displayed exceptionally high presence/absence variation of MAX effectors (average Hamming distance between pairs of isolates: 0.123 and 0.095; Figure 4D), but only limited sequence variation at single-copy core MAX effectors (average Hamming distance between pairs of isolates: 0.017 and 0.012; Mann-Whitney U-tests, p<0.05; Figure 4C).

### Loss of MAX effectors in specific lineages does not appear to be associated with host specificity

The comparison of the MAX effector content in the genomes of 120 *P. oryzae* isolates revealed extensive presence/absence polymorphism between host-specific groups (S3 Table). To address the underlying evolutionary mechanisms, we tested experimentally the hypothesis that MAX effector losses are massively related to escape from receptor-mediated non-host resistance. Indeed, the loss of MAX effectors in specific lineages of *P. oryzae* could primarily serve to escape from non-host resistance during infections of novel plant species carrying immune receptors specifically recognizing these effectors. To test this hypothesis, we focused on the *Oryza-* and *Setaria-infecting* lineages, as previous investigations suggested that the Oryza-infecting lineage emerged by a host shift from *Setaria* and we found both groups to be closely related (Figure 4) [21, 42]. Our strategy was to introduce into the Oryza-isolate Guy11 MAX effectors absent from the *Oryza* lineage but present in the *Setaria* lineage, and to assess the ability of these transgenic isolates to infect rice.

We identified three MAX orthogroups that were largely or completely absent from the *Oryza* lineage, but present in the majority of isolates of the other lineages (S3 Table). Orthogroup MAX79 (OG0011591-1) was absent in all 52 Oryza-infecting isolates, while MAX83 (OG0011907), and MAX89 (OG0012141) were absent in 50 and 46 of them, respectively (S3 Table). Constructs carrying the genomic sequence of *MAX79, MAX83 or MAX89* derived from the *Setaria* isolate US0071 and under the control of the strong infection specific promoter of the effector *AVR-Pia* were generated and stably introduced into Guy11. For each construct, three independent transgenic lines were selected. Transgene insertion was verified by PCR and the expression of transgenes was measured by qRT-PCR (S7 Figure). To test whether the selected MAX effectors trigger immunity in rice, the transgenic isolates were spray-inoculated onto a panel of 22 cultivars representative of the worldwide diversity of rice (S4 Table).

As controls, we used the MAX effectors *AVR-Pia*, which is rare outside the *Oryza* and *Setaria* lineages, and *AVR1-CO39*, which is absent or pseudogenized in the *Oryza* lineage, but present in all other host-specific lineages including *Setaria*. Both effectors are detected in rice by the paired NLR immune receptors RGA4 and RGA5 from the *Pi-a/Pi-CO39* locus and thereby contribute, respectively, to host or non-host resistance in this plant species [43, 44].

As expected, isolates expressing *AVR1-CO39* or *AVR-Pia* triggered resistance in the rice variety Aichi Asahi that carries *Pi-a*, but caused disease on Nipponbare (*pi-a*^*-*^) and other varieties lacking this *R* locus (Figure 5; S4 Table). Unlike the positive controls, the effectors MAX79, MAX83 and MAX89 were not recognized and did not induce resistance in any of the tested rice cultivars (Figure 5; S4 Table). The disease symptoms caused by the transgenic isolates carrying these effectors were similar to those observed for wild-type Guy11 or Guy11 isolates carrying an *RFP* (red fluorescent protein) construct. This suggests that these effectors do not significantly increase the virulence of Guy11.

**Figure 5.**
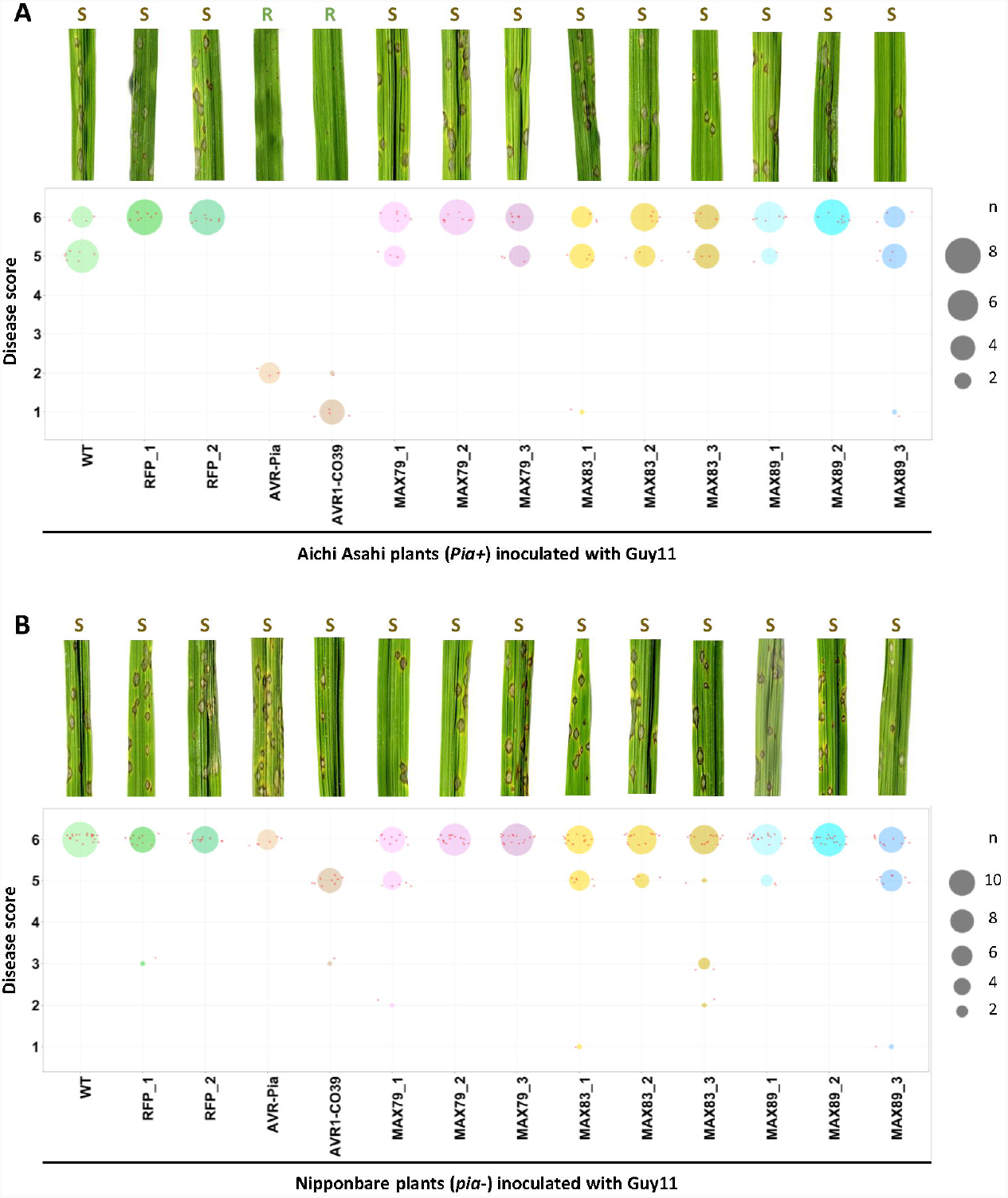
AVR1-CO39 contributes to non-host specificity in rice but not MAX79, MAX83 or MAX89. Wild type and transgenic isolates of *P. oryzae* Guy11 expressing the *RFP (red fluorescent protein), A VR-Pia, A VR1-CO39, MAX79, MAX83* or *MAX89gene* were spray-inoculated at 40 000 spores/ml on three-week-old rice plants of the cultivars Aichi Asahi (A) and Nipponbare (B). For each condition, representative disease phenotypes on rice leaves at seven days post-inoculation are shown (top panels, R: resistance, S: susceptibility). Disease phenotypes were also scored (from 1 [complete resistance] to 6 [high susceptibility]) on leaves from three to five individual rice plants and data are shown as dot plots (bottom panels). The size of each circle is proportional to the number of replicates (n) matching the corresponding score for each condition. Small red dots correspond to individual measurements. The experiment was performed twice for Aichi Asahi and four times for Nipponbare for all isolates except for WT, AVR-Pia, and AVR1-CO39 control isolates. For these isolates, experiments were performed once on Aichi Asahi and twice on Nipponbare because disease phenotypes are well characterized in the literature.

These experiments show that despite their loss in the Oryza-infecting lineage of *P. oryzae*, and unlike AVR1-CO39, the effectors MAX79, MAX83, and MAX89 do not seem to induce non-host resistance in rice. Consequently, other mechanisms than escape from host immunity contributed to the loss of these MAX effectors during the putative host shift of *P. oryzae* from *Setaria to Oryza*.

### MAX effectors display signatures of balancing selection

To investigate the impact of balancing selection on MAX effector evolution, we focused on singlecopy core, softcore, and shell orthogroups to avoid the possible effect of gene paralogy. We then computed *π* (nucleotide diversity per bp), *F*_*ST*_ (the amount of differentiation among lineages [45]), *π* _N_ (non-synonymous nucleotide diversity), *π*_S_ (synonymous nucleotide diversity), and π_N_/π_S_ (the ratio of non-synonymous to synonymous nucleotide diversity). Large values of *π* and π_N_/π_S_, in particular, are possible indicators of a gene being under balancing selection.

Nucleotide diversity (π) differed significantly between groups of genes (Kruskal-Wallis test, H=509.9, d.f.=2, *p*<0.001; Figure 6A; S5 Table). *π* was significantly higher for the set of MAX effectors (average *π*. 0.0104, standard deviation: 0.0137), than for other secreted proteins (average *π*. 0.0079, standard deviation: 0.020), and other genes (average *π*. 0.0049, standard deviation: 0.014; Mann-Whitney U-tests, *p<*0.05), showing that MAX effectors, and to a smaller extent other secreted proteins, are more variable than a typical gene. At the lineage level, however, nucleotide diversity at MAX effectors tended to not significantly differ from other putative effectors, or other genes (S5 Table).

**Figure 6.**
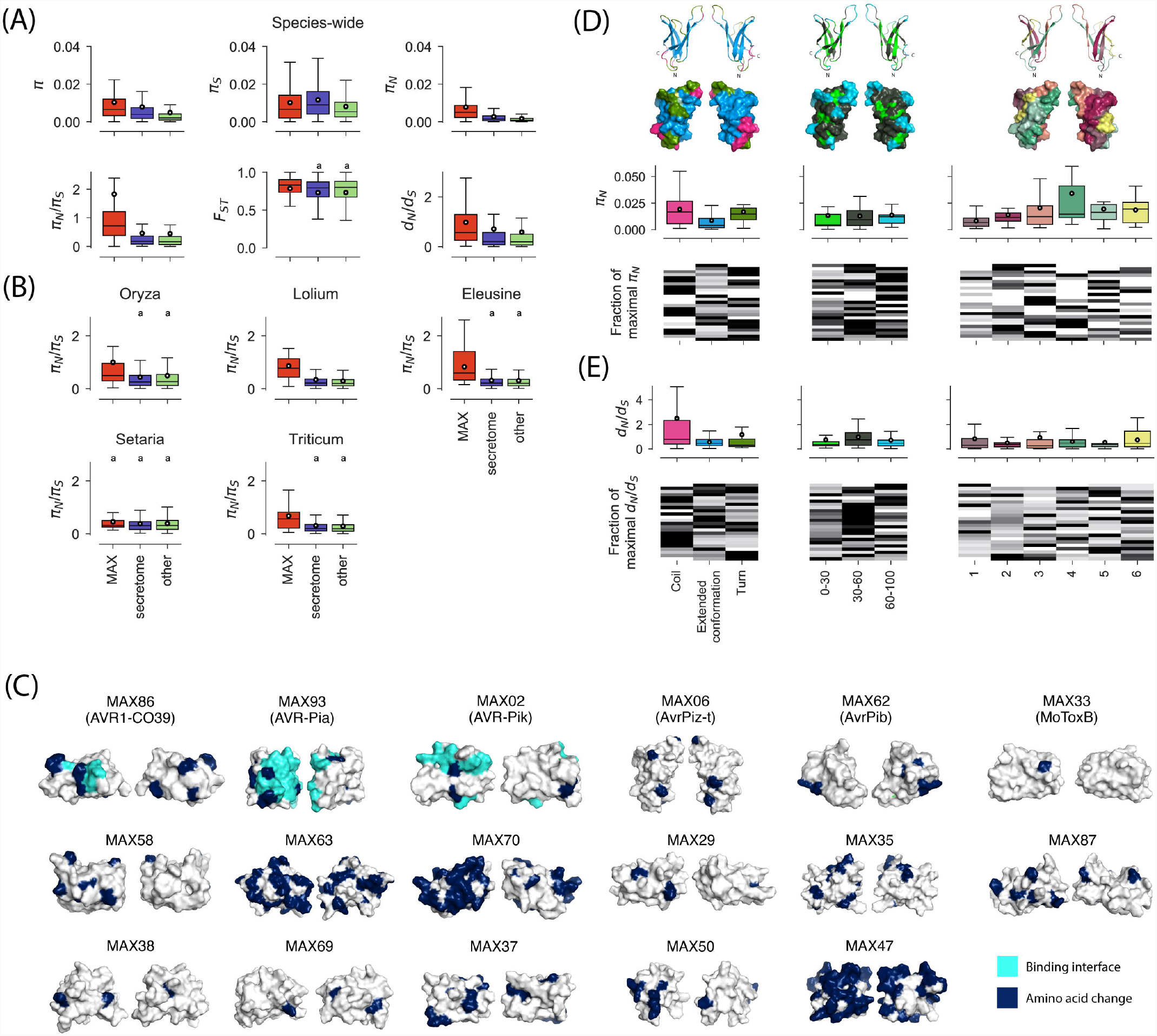
Summary statistics of polymorphism and divergence at MAX effectors, other secreted proteins (i.e., secretóme), and other genes of *P. oryzae*. (A) Species-wide estimates of *it* (nucleotide diversity per bp), *F*_*ST*_ (the amount of differentiation among lineages), *π*_*N*_ (non-synonymous nucleotide diversity per bp), *π*_*S*_ (synonymous nucleotide diversity ), *π*_*N*_*/π*_*S*_ (the ratio of non-synonymous to synonymous nucleotide diversity), *dN/ds* (the ratio of non-synonymous to synonymous rates of substitutions). (B) Lineage-specific estimates of *π_N_/π_S_-*(C) Amino acid changes segregating in *P. oryzae* at MAX effectors with an avirulence function and MoToxB (first row) and MAX effectors with *π_N_/π_S_>2* (next rows); amino acid changes are shown in dark blue and known binding interfaces in light blue; all effectors are represented with the same orientation as AvrPiz-t in panel (B). Note that the *π*_*N*_*/π*_*S*_ ratio is >2 for the avirulence genes *AvrPiz-t* and AVR-Pik. Proteins are displayed twice in panels C and E, with one copy rotated 180 degrees around a vertical axis. The interface involved in binding with host proteins is known for AVR1-CO39, AVR-Pia, and AVR-Pik only ([50-53]). (D) and (E) Species-wide estimates of *π*_*N*_ and *dN/ds* computed at MAX effectors with signatures of balancing selection *(π*_*N*_*/π*_*S*_*>*1; panel D) and signatures of directional selection (*π*_*N*_*/π*_*S*_;1 panel E) for different classes of structural features highlighted on the three-dimensional structure of *AvrPiz-t* above panel D: (i) secondary structure elements, with three subclasses: “coils”, “extended conformation”, and “turns”; (ii) solvent accessibility percentage of the Van der Waals surface of the amino acid side chain, with three sub-classes: 0-30% (buried), 30-60% (intermediate), and 60-100% (exposed); (iii) structural domains, with six subclasses that grouped the coil, extended conformation and turn residues that define the six beta strands characteristic of MAX effectors. In the heatmaps, each line represents a MAX effector. For a given MAX effector and a given structural feature, the darkest color indicates the class of the structural feature for which the summary statistic is the highest. Only single-copy core, softcore, and shell groups of orthologous genes were included in the calculations. Shared superscripts indicate non-significant differences (post-hoc Mann-Whitney U-tests, *p*>0.05). A number of data points were cropped from plots in (A) and (B) for visually optimal presentation but included in statistical tests. In box plots, the black circle is the mean, the black line is the median.

In addition to having greater nucleotide variation than other genes at the species level, MAX effectors also displayed a higher ratio of non-synonymous to synonymous nucleotide diversity (Figure 6B; S5 Table). The *π*_*N*_*/π*_*S*_ ratio differed significantly between groups of genes (Kruskal-Wallis tests *H*=101.4, *d.f.=2*. *p<*0.001), and the excess of non-synonymous diversity was significantly, and markedly, higher for MAX effectors (average *π*_*N*_*/π*_*S*_; 1.826, standard deviation: 3.847) than for other effectors (average *π*_*N*_*/*π_*S*_: 0.461, standard deviation: 1.600), and other genes (average *π*_*N*_/*π*_*S*_ 0.448, standard deviation: 1.463; Mann-Whitney U-tests, *p*< 0.05). The higher *π*_*N*_/*π*_*S*_ MAX effectors was mostly driven by differences in *π*_*N*_ (Figure 6A; S5 Table). Twenty MAX effectors displayed values in the top 5% percentile of non-effector genes, far exceeding the four genes expected by chance (*p*<0.05). More specifically, 26 MAX effectors displayed *π*_*N*_*/π*_*S*_ values greater than 1, which is the value expected under neutrality. This included three well-known avirulence genes: *AVR1-CO39* (*π*_*N*_/*π*_*S*_=2.564), *AVR-Pik* (*π*_*N*_/*π*_*S*_=15.574), and *AvrPiz-t* (*π*_*N*_/*π*_*S*_=1.431). The average *π*_*N*_*/π*_*S*_ ratio was also higher at MAX effectors than other secreted proteins and other genes in all lineages, with significant differences in four lineages (Mann-Whitney U-tests, *p<*0.05), and the average *π*_*N*_/*π*_*S*_ was greater than one in the *Oryza* lineage (Figure 6B; S5 Table). Seven to eleven MAX effectors had *π*_*N*_*/π*_*S*_*>1* at the lineage level, representing 8% (*Setaria*-infecting lineage) to 41% (*Lolium* infecting lineage) of MAX effectors with a defined *π*_*N*_*/π*_*S*_ ratio (S2 Data).

*π*_*N*_/*π*_*S*_ >l is a strong indication of multiallelic balancing selection (*i e*., multiple alleles at multiple sites are balanced), as single sites under very strong balancing selection cannot contribute enough non-synonymous variability to push the *π*_*N*_*/π*_*S*_ ratio above one [46]. To assess whether the adaptation of lineages to their respective hosts may contribute to the species-wide excess of non-synonymous diversity detected at MAX effectors, we estimated population differentiation. The differentiation statistic *F*_*ST*_ differed significantly between groups of genes (Kruskal-Wallis tests *H*=8.731, *d.f.=2*. *p*=0.013), and differentiation was significantly higher for MAX effectors than for other secreted proteins and other genes (Figure 6A; S5 Table). *F_ST_* was also significantly, albeit relatively weakly, correlated with *π*_*N*_*/π*_*S*_ at MAX effectors (Spearman’s *p*. 0.304, *p*=0.007; S8 Figure). These observations indicate that between-lineages differences in allele frequencies are greater for MAX effectors than for other secreted proteins or other genes, which may result from divergent selection exerted by hosts.

### MAX effectors display signatures of recurrent directional selection

To detect adaptive molecular evolution, we collected orthologous sequences from outgroup *Pyricularia sp*. LS [47, 48] and estimated the *d*_*N*_/*d*_*S*_ ratio (the ratio of non-synonymous to synonymous substitution rates) using a maximum likelihood method [49]. Note that *d*_*N*_/*d*_*S*_ was not computed at the intra-specific level, but computed along the branches connecting the outgroup and isolates from the ingroup. Outgroup sequences could be retrieved for 10,174 out of 14,664 single-copy orthogroups, including 66 out of 94 single-copy orthologs of MAX effectors. The *d*_*N*_*/d*_*S*_ ratio differed significantly between groups of genes (Kruskal-Walli. s tests 27=45.812, *d.f.=2*. *p<*0.001; Figure 6A; S5 Table), and was higher for MAX effectors (average *d*_*N*_/*d*_*S*_. 0.977, s.d.: 1.316) than for other secreted proteins (average *d*_*N*_/*d*_*S*_ 0.711, s.d.: 1.722), and other genes (average *d*_*N*_/*d*_*S*_. 0.584, s.d.: 1.584; Mann-Whitney U-tests, *p<*0.05). The same pattern of higher *d*_*N*_/*d*_*S*_ for MAX effectors was observed at the lineage level (S5 Table). Twenty-four of the 66 MAX effectors with outgroup sequence (i.e., 36.4%) showed *d*_*N*_/*ds >l* (S2 Data), which is a strong indication of directional selection, *d*_*N*_/*d*_*S*_ > 1 is only expected for genes that have experienced repeated bouts of directional selection which led to repeated fixations of amino-acid substitutions [46]. Eleven MAX effectors displayed signatures of both multiallelic balancing selection (*π*_*N*_/*π*_*S*_ > 1) and multiallelic directional selection (*d*_*N*_/*d*_*S*_ > 1).

The divergence data, therefore, indicate that a scenario of molecular co-evolution involving repeated selective sweeps may apply to a substantial fraction (at least one-third) of MAX effectors.

### Structural determinants of polymorphism and divergence at MAX effectors

Different parts of proteins can be under different selective forces. To investigate if this is the case in MAX effectors, we examined the relationship between three different measures of diversity and three structural features. The measures of diversity were (1) the probability of an amino acid being polymorphic, (2) the non-synonymous nucleotide diversity *π*_*N*_, and (3) the *d*_*N*_/*d*_*S*_ ratio. The analyzed structural features were: (1) secondary structure annotations with the three subclasses “extended conformation”, “coils”, and “turns”; (2) solvent accessibility percentage of the Van der Waals surface of the amino acid side chain, with the three sub-classes: 0-30% (buried), 30-60% (intermediate), and 60-100% (exposed); (3) structural domains, with six subclasses that grouped the coil, extended conformation and turn residues that define the six beta strands characteristic of MAX effectors. Structural features were determined using MAX effector structures predicted by homology modeling and computing with STRIDE.

For the relationship between the probability of residues being polymorphic and the structural features, we used a generalized linear mixed model with a set of predictor variables. The fixed effects were the structural features and the model was fitted using maximum likelihood estimation, with MAX effector modeled as a random effect. Only a single structural feature, the solvent accessibility had a statistically significant effect on the probability of amino acid polymorphism (S1 Text). Predicted probabilities of amino acid change were higher for accessibility class 60-100% (95% prediction interval: 0.1294-0.1295), than for accessibility classes 30-60% (95% prediction interval: 0.1029-0.1049) and 0-30% (95% prediction interval: 0.0834-0.0835). A major factor explaining the variability of the response variable turned out to be the identity of MAX effectors. Indeed, the random effect (σ: 1.17) had a larger standard deviation than the largest fixed-effect factor (coefficient for accessibility class 60-100%: 0.64) (S1 Text).

To visualize the localization of polymorphisms, we projected the distribution of amino acid changes on the surface of protein structure models of two types of MAX effectors: those with an avirulence function and MAX effectors with the strongest signatures of multiallelic balancing selection (*π*_*N*_*/π*_*S*_>2; Figure 6C; S9 Figure). For the three effectors whose binding interfaces have been experimentally characterized (AVR1-CO39, AVR-Pia, AVR-Pik [50-53]), a substantial fraction of amino acid changes co-localized with residues interacting with immune receptors and, presumably, also with their host target proteins. Polymorphic residues are, therefore, potentially good predictors for binding interfaces in MAX effectors and the specific surface regions, where polymorphisms cluster in several MAX effectors (such as MoToxB, MAX58, MAX87, MAX69, or MAX50) could correspond to interfaces that bind immune receptors and/or host target proteins (Figure 6C).

To determine which parts of the MAX structure is most responsible for the high level of standing variation in these effectors, we calculated for the three different structural features and their subclasses the non-synonymous nucleotide diversity *π*_*N*_ (S10 Figure). We restricted this analysis to the 25 MAX effectors exhibiting balancing selection (*π*_*N*_*/π*_*S*_> 1 ) and we used *π*_*N*_ and not *π*_*N*_*/π*_*S*_ because the latter tended to be undefined due to relatively short sequence lengths. Non-synonymous nucleotide diversity *π*_*N*_ differed between subclasses (Kruskal-Wallis tests *H*=8.504, *d.f.=2*. *p*=0.014; Figure 6D; S6 Table) and was higher at coils and turns, than at extended conformations (coils: *π*_*N*_=0.0191; turns: *π*_*N*_ = 0.0167; extended conformations: *π*_*N*_ = 0.0086; posthoc Mann-Whitney U-tests, *p<*0.05). Ten and nine MAX effectors displayed their highest values of *π*_*N*_ in coils and turns, respectively, *π*_*N*_ did not significantly differ between relative solvent accessibility subclasses (Kruskal-Wallis tests *H*=2.308, *d.f.=2*. *p*=0.315), but differences were marginally significant between structural domains (Kruskal-Wallis tests *H*=11.035, *d.f.=5*, *p*=0.051). The third, fourth, and fifth beta strands displayed the highest levels of non-synonymous diversity *(π*_*N*_*=* 0.0341, *π*_*N*_ =0.0341, and *π*_*N*_ =0.0341, respectively), and 18 out of 25 MAX effectors displayed their highest values of *π*_*N*_ at one of these three beta strands (S6 Table).

To identify the parts of MAX effector structures that experience multiallelic directional selection, we analyzed the 23 proteins with *d*_*N*_/*d*_*S*_ >1 This showed that differences in *d*_*N*_/*d*_*S*_ were most pronounced between subclasses of structural features (Kruskal-Wallis tests *H=5*.499, *d.f.=2*. *p*=0.064), with higher average *d*_*N*_/*d*_*S*_ values for coils and turns (*d*_*N*_/*d*_*S*_ =2.490 and *d*_*N*_/*d*_*S*_ =1.184, respectively) than extended conformations (*d*_*N*_/*d*_*S*_ =0.573) (Figure 6E; S6 Table). The average *d*_*N*_/*d*_*S*_ was also close to one for the 30-60% subclass of relative solvent accessibility (*d*_*N*_/*d*_*S*_ =0.994), and 12 MAX effectors with signatures of directional selection had their highest *d*_*N*_/*d*_*S*_ values for this subclass, although differences were not significant.

Overall, these analyses show that multiallelic balancing and directional selection acted preferentially on coils and turns, but that the impact of two forms of selection on structural domains and solvent accessibility subclasses differs.

## DISCUSSION

### MAX effectors as model systems to investigate effector evolution

Effectors involved in coevolutionary interactions with host-derived molecules are expected to undergo non-neutral evolution. Yet, the role of natural selection in shaping polymorphism and divergence at effectors has remained largely elusive [2]. Despite the prediction of large and molecularly diversified repertoires of effector genes in many fungal genomes, attempts to probe into the evolutionary drivers of effector diversification in plant pathogenic fungi have been hindered by the fact that, until recently, no large effector families had been identified. In this study, we overcome the methodological and conceptual barrier imposed by effector hyperdiversity by building on our previous discovery [17] of an important, structurally-similar, but sequence-diverse family of fungal effectors called MAX. We used a combination of structural modeling, evolutionary analyses, and molecular plant pathology experiments to provide a comprehensive overview of polymorphism, divergence, gene expression, and presence/absence at MAX effectors. When analyzed species-wide or at the level of sub-specific lineages, ratios of non-synonymous to synonymous nucleotide diversity, as well as ratios of non-synonymous to synonymous substitutions, were consistently higher at MAX effectors than at other loci. At the species level, the two ratios were also significantly higher than expected under the standard neutral model for a large fraction of MAX effectors. The signatures of adaptive evolution detected at MAX effectors, combined with their extensive presence/absence variation, are consistent with their central role in coevolutionary interactions with host-derived ligands that impose strong selection on virulence effectors.

### Adaptive evolution of MAX effectors

Rates of evolution determined from orthologous comparisons with outgroup sequences revealed that, for a large fraction of MAX effectors, non-synonymous changes have accumulated faster than synonymous changes. The fast rate of amino-acid change at MAX effectors is consistent with a classic arms race scenario, which entails a series of selective sweeps as new virulent haplotypes -capable of avoiding recognition by plant immune receptors that previously prevented pathogen multiplication -spread to high frequency [54, 55]. Furthermore, it is important to note that although large values of the *d*_*N*_/*d*_*S*_ ratio provide strong evidence for directional selection, small values do not necessarily indicate the lack thereof, as *d*_*N*_/*d*_*S*_ ratios represent the integration of genetic drift, constraint, and adaptive evolution [50][56]. Much of the adaptive changes at MAX effectors probably took place before the radiation of *P. oryzae* on its various hosts. However, the observation that *d*_*N*_/*d*_*S*_ values determined from orthologous comparisons with outgroup are higher at the species level than at the sub-specific lineage level indicates that part of the signal of directional selection derives from inter-lineage amino acid differences associated with host-specialization. Our structural modeling indicates that it is preferentially “turns” and “coils”, but also residues with intermediate solvent accessibility, which often evolve at an unusually fast rate, and therefore that these are probably the residues of MAX proteins preferentially involved in coevolutionary interactions with host-derived molecules.

MAX effectors are characterized by a remarkable excess of non-synonymous polymorphism, compared to synonymous polymorphism, at the species level, but also -albeit to a lesser extent -at the sub-specific lineage level. This raises the question of how polymorphisms are maintained in the face of adaptive evolution, given that selective sweeps under a classic arms race scenario are expected to erase variation [6, 9]. Directional selection restricted to host-specific lineages -i.e., local adaptation -may contribute to the signature of multiallelic balancing selection observed at the species level. The observation of a positive correlation between *π*_*N*_*/π*_*S*_ and the differentiation statistic *F*_*ST*_, together with the fact that most MAX effectors are monomorphic at the lineage level, are consistent with a role of divergent selection exerted by hosts in the maintenance of species-wide diversity at MAX effectors. However, the finding that MAX effectors with a defined *π*_*N*_*/π*_*S*_ at the sub-specific lineage level (i.e., MAX effectors with *π*_*S*_ ≠ 0) present a higher ratio than the other genes also indicates that the adaptive evolution process is not simply one of successive selective sweeps. This is consistent with balancing selection acting at the lineage level, through which polymorphisms in MAX virulence effectors are maintained due to spatiotemporal variation in selection pressures posed by the hosts - a process known as the trench-warfare model [54]. MAX effectors can experience varying selection pressures due to differences in both arsenals of immune receptors and repertoires of virulence targets across host populations. This means that the variability of effectors can result both from their evolution to avoid detection, and from their evolution to maintain their virulence activity *(e.g*., by targeting one or more potentially polymorphic host proteins to suppress avirulence or basal immunity, or to manipulate other host cellular processes). Our structural modeling suggests in particular that the “coils” and “turns” are the preferred substrate of these coevolutionary interactions leading to the maintenance of elevated polymorphism at MAX virulence effectors. Mirroring the existence of hypervariable MAX effectors, we also detect a substantial proportion of effectors that show no variability, either at the species level or at the lineage level. However, the lack of variability does not necessarily mean they have no impact on virulence. It is possible that their role in virulence is associated with evolutionary constraints that restrict their variability to a limited region of sequence space. Core, monomorphic MAX effectors could be prime targets for genetically-engineered NLRs [57].

### Expression kinetics of MAX effectors

Expression profiling showed that the MAX effector repertoire was induced specifically and massively during infection. Depending on the host genotype, between 64 and 78% of the MAX effectors were expressed and expression was particularly strong during the early stages of infection. These findings are consistent with previous studies that analyzed genome-wide gene expression during rice infection or specifically addressed MAX effector expression, and they reinforce the hypothesis that MAX effectors are crucial for fungal virulence and specifically involved in the biotrophic phase of infection [17, 38].

How this coordinated deployment of the MAX effectors is regulated remains largely unknown. Genome organization does not seem to be a major factor, since MAX effectors do not colocalize and more generally, there is no clustering of effectors in the *P. oryzae* genome, only a slight enrichment in subtelomeric regions [38, 58]. This differs from other pathogenic fungi, such as *Leptosphaeria maculans*, for which early-expressed effectors are clustered in AT-rich isochores, and co-regulated by epigenetic mechanisms [59]. Analysis of promoter regions of MAX effectors did not identify common DNA motifs that may be targeted by transcription factors, and no such transcriptional regulators that would directly regulate large fractions of the effector complement of *P. oryzae* have been identified yet. The few known transcriptional networks controlled by regulators of *P. oryzae* pathogenicity generally comprise different classes of fungal virulence genes, such as secondary metabolism genes or carbohydrate-active enzymes; they are not restricted to effectors. Recently, it was shown that Rgs 1, a regulator of G-protein signaling necessary for appressorium development, represses the expression of 60 temporally co-regulated effectors in axenic culture and during the pre-penetration stage of plant infection [60]. Of these, six belong to the MAX family and their expression is affected in *cer7* and Δ*rgs1* mutants: MGG_1004T0 [15, 38]. MGG_15443T0 [16, 38]. MGG_08817T0 [15]. MGG_17266T0 [15-17, 38] and MAX15 (MGG_05424T0) and MAX67 (MGG_16175T0), both identified in our study. This represents only 5% of the MAX effectors predicted to date (S2 Table) and suggests that multiple complementary mechanisms contribute to the precise coordination of MAX effector expression during rice invasion.

Expression profiling also revealed that the plant host genotype strongly influenced the expression of the MAX effector repertoire, suggesting that plasticity in effector expression may contribute to the adaptation of *P. oryzae* to its hosts. MAX effectors were stronger expressed in the more resistant Kitaake rice variety than in highly susceptible Maratelli rice. This is reminiscent of other pathogenic fungi, such as *Fusarium graminearum* and *L. maculans*, for which a relationship between host resistance levels and effector expression was established [61, 62]. An expression analysis of MAX effectors in isolates infecting a wider range of host plants with varying resistance levels could be conducted to further investigate the connection between plant resistance and MAX effectors’ expression.

### Presence/absence polymorphism of MAX effectors

Pangenome analyses demonstrated extensive variability in the MAX effector repertoire. In cases where MAX effectors are specifically absent from some lineages, but present in most or all others, it is tempting to hypothesize that they experienced immune-escape loss-of-function mutations that directly contributed to host range expansion or host shifts. A possible example of such a mechanism is the non-MAX effector *PWT3* of *P. oryzae* that is specifically absent from the *Triticum*-infecting lineage [29]. *PWT3* triggers resistance in wheat cultivars possessing the *RWT3* resistance gene [63]. and its loss was shown to coincide with the widespread deployment of *RWT3* wheat. Similarly, the loss of the effector *AVR1-C039 (MAX86)*, which is specifically absent from the *Oryza*-infecting lineage and that is detected by the rice NLR immune receptor complex RGA4/RGA5, has been suggested to have contributed to the initial colonization of rice by the *Setaria*-infecting lineage [20, 31, 64]. Two other orthologous *P. oryzae* effectors, *PWL1* and *PWL2*, exclude Eleusine and rice-associated isolates from infecting *Eragrostis curvula*, and can, therefore, also be considered as host-specificity determinants [28, 65]. Interestingly, ALPHAFOLD predicts PWL2 to adopt a MAX effector fold [66]. In our study, however, gene knock-in experiments with *MAX79, MAX83*, and *MAX89 -*specifically absent from the (*Oryza*-infecting lineage -did not reveal a strong effect on virulence towards a large panel of rice varieties. Hence, unlike AVR1-CO39, these effectors are not key determinants of host-specificity. This suggests that overcoming non-host resistance is not the only and maybe not the main evolutionary scenario behind the specific loss of MAX effectors in the *Oryza*-infecting lineage. A possible alternative mechanism that can explain massive MAX effector loss during host shifts is a lack of functionality in the novel host. Some MAX effectors from a given lineage may have no function in the novel host, simply because their molecular targets are absent or too divergent in the novel host. Cellular targets of fungal effectors remain unknown for the most part, but knowledge of the molecular interactors of MAX effectors may help shed light on the drivers of their presence/absence polymorphism.

### Concluding remarks

The discovery of large, structurally-similar, effector families in pathogenic fungi and the increasing availability of high-quality whole genome assemblies and high-confidence annotation tools, pave the way for in-depth investigations of the evolution of fungal effectors by interdisciplinary approaches combining state-of-the-art population genomics, protein structure analysis, and functional approaches. Our study on MAX effectors in the model fungus and infamous cereal killer *P. oryzae* demonstrates the power of such an approach. Our investigations reveal the fundamental role of directional and balancing selection in shaping the diversity of MAX effector genes and pinpoint specific positions in the proteins that are targeted by these evolutionary forces. This type of knowledge is still very limited on plant pathogens, and there are very few studies compared to the plethoric literature on the evolution of virulence factors in human pathogens. Moreover, by revealing the concerted and plastic deployment of the MAX effector repertoire, our study highlights the current lack of knowledge on the regulation of these processes. A major challenge will now be to identify the regulators, target proteins and mode of action of MAX effectors, in order to achieve a detailed understanding of the relationships between the structure, function and evolution of these proteins.

## METHODS

### Genome assemblies, gene prediction, and pan-genome analyses

Among the 120 genome assemblies included in our study, 66 were already assembled and publicly available, and 54 were newly assembled (S1 Table). For the 54 newly assembled genomes, reads were publicly available for 50 isolates, and four additional isolates were sequenced (available under BioProject PRJEB47684). For the four sequenced isolates, DNA was extracted using the same protocol as in ref. [67]. TruSeq nano kits were used to prepare DNA libraries with insert size of ∼500bp for 150 nucleotide paired-end indexed sequencing with Illumina HiSeq 3000. For the 54 newly generated assemblies, CUTADAPT [68] was used for trimming and removing low-quality reads, reads were assembled with ABYSS 1.9.0 [69] using eight different K-mer sizes, and we chose the assembly produced with the K-mer size that yielded the largest N50. For all 120 genome assemblies, genes were predicted by BRAKER 1 [70] using RNAseq data from ref. [21] and protein sequences of isolate 70-15 (Ensembl Fungi release 43). To complement predictions from Braker, we also predicted genes using Augustus 3.4.0 [56] with RNAseq data from ref. [21]. protein sequences of isolate 70-15 (Ensembl Fungi release 43), and *Magnaporthe grisea* as the training set. Gene predictions from Braker and Augustus were merged by removing the genes predicted by AUGUSTUS that overlapped with genes predicted by Braker. Repeated elements were masked with RepeatMasker 4.1.0 (http://www.repeatmasker.org/). The quality of genome assembly and gene prediction was checked using BUSCO 4.0.4 [32]. The homology relationships among predicted genes were identified using ORTHOFINDER V2.4.0 [36]. The size of pan- and core-genomes was estimated using rarefaction, by resampling combinations of one to 119 genomes, taking a maximum of 100 resamples by pseudo-sample size. Sequences for each orthogroup were aligned at the codon level (i.e., keeping sequences in coding reading frame) with Translatorx 1.1 [71]. using MAFFT v7 [72] as the aligner and default parameters for Gblocks 0.91B [73]. The effect of assembly properties, host of origin, and study of origin on the number of predicted genes computed from the orthology table was analyzed in Python 3.7 using the function in Pearsonr in Scipy.stats 1.10.1, and functions Formula.api.ols and Stats.anova.anova_lm in Statsmodels 0.15.0.

### Identification of effectors sensu lato, and MAX effectors

We predicted the secretome by running Signal P 4.1 [74]. TargetP 1.1 [75]. and Phobius 1.01 [62] to identify signal peptides in the translated coding sequences of 12000 orthogroups. Only proteins predicted to be secreted by at least two methods were retained. Transmembrane domains were identified using TMHMM [76] and proteins with a transmembrane domain outside the 30 first amino acids were excluded from the predicted secretome. Endoplasmic reticulum proteins were identified with PS-SCAN (https://ftp.expasy.org/databases/prosite/ps_scan/), and excluded.

To identify MAX effectors, we used the same approach as in the original study that described MAX effectors [17]. We first used PSI-BLAST 2.6.0 [33] to search for homologs of known MAX effectors (AVR1-CO39, AVR-Pia, AvrPiz-t, AVR-PikD, and ToxB) in the predicted secretome. Significant PSI-BLAST hits (e-value < e-4) were aligned using a structural alignment procedure implemented in TM-ALIGN [35]. Three rounds of HMMER [34] searches were then carried out, each round consisting of alignment using TM-ALIGN version 20140601, model building using HMMBUILD, and HMM search using HMMSEARCH (e-value < e-3). Only proteins with two expected conserved cysteines less than 33-48 amino acids apart were retained in the first two rounds of HMMER searches, as described in ref. [17].

Subsequent evolutionary analyses were conducted on three sets of orthogroups: MAX effectors, putative effectors, and other genes. The “MAX” group corresponded to 80 orthogroups for which at least 10% of sequences were identified as MAX effectors. The “secreted proteins” groups corresponded to 3283 orthogroups that were not included in the MAX group, and for which at least 10% of sequences were predicted to be secreted proteins. The last group included the remaining 11404 orthogroups.

For missing MAX effector sequences, we conducted an additional similarity search to correct for gene prediction errors. For a given MAX orthogroup and a given isolate, if a MAX effector was missing, we used BLAST-N to search for significant hits using the longest sequence of the orthogroup as the query sequence, and the isolate’s genome assembly as the subject sequence (S3 Table). We also corrected annotation errors, such as the presence of very short (typically <50bp) or very long (typically >500bp) introns, missing terminal exons associated with premature stops, or frameshifts caused by indels. All these annotation errors were checked, and corrected manually if needed, using the RNAseq data used in gene prediction in the Integrative Genome viewer [77, 78]. We also found that some orthogroups included chimeric genes resulting from the erroneous merging of two genes that were adjacent in assemblies. This was the case for orthogroups OG0000093 and OG0010985, and we used RNA-seq data in the Integrative Genome Viewer to split the merged genic sequences and keep only the sequence corresponding to a MAX effector.

For evolutionary analyses conducted on single-copy orthologs, the 11 orthogroups that included paralogous copies of MAX effectors were split into sets of orthologous sequences using genealogies inferred using RAxML v8 [37]. yielding a total of 94 single-copy MAX orthologs, of which 90 orthologs passed our filters on length and sample size to be included in evolutionary analyses (see below). For each split orthogroup, sets of orthologous sequences were assigned a number that was added to the orthogroup’s identifier as a suffix (for instance paralogous sequences of orthogroup OG0000244 were split into orthogroups OG0000244_l and OG0000244_2). Sequences were re-aligned using TRANSLATORX (see above) after spliting orthogroups.

All genome assemblies, gene models, aligned coding sequences for all orthogroups, and single-copy orthologs, are available in Zenodo, doi: 10.5281/zenodo.7689273 and doi: 10.5281/zenodo.8052494.

### Analysis of population subdivision

Population structure was inferred from SNPs identified in GBLOCKS-cleaned alignments of coding sequences at 7317 single-copy core orthologs (described in section *Genome assemblies, gene prediction, and pan-genome analyses*). We kept only one randomly chosen four-fold degenerate synonymous site per single-copy core ortholog. We used the SNMF method from the LEA package in R [40] to infer individual ancestry coefficients in *K* ancestral populations. We used SPLITSTREE version 5.3 [41] to visualize relationships between genotypes in a phylogenetic network, with reticulations to represent the conflicting phylogenetic signals caused by homoplasy.

### Homology modeling of MAX effectors

To check that orthogroups predicted to be MAX effectors had the typical 3D structure of MAX effectors with two beta sheets of three beta strands each, eight experimental structures with MAX-like folds were selected as 3D templates for homology modeling (PDB identifiers of the templates: 6R5J, 2MM0, 2MM2, 2MYW, 2LW6, 5A6W, 5Z1V, 5ZNG). For each of the 94 MAX orthologous groups, one representative protein was selected and homology models of this 1D query relative to each 3D template were built using Modeller [79] with many alternative query template threading alignments. The structural models generated using the alternative alignments were evaluated using a combination of six structural scores (DFIRE [80]. GOAP [81]. and QMEAN’s E_1D_, E_2D_, E_3D_ scores [82]). A detailed description of the homology modeling procedure is provided in S2 Text The best structural models for the 94 representative sequences of each group of MAX orthologs are available at https://pat.cbs.cnrs.fr/magmax/model/. The correspondence between MAX orthogroups identifiers used in homology modeling and MAX orthogroups identifiers resulting from gene prediction is given in S2 Table. Protein models were visualized with PYMOL 2.5 [83].

### Evolutionary analyses

Lineage-level analyses were conducted on a dataset from which divergent or introgressed isolates were removed (G17 from *Eragrostis*, Bm88324 & Bd8401 from *Setaria*, 87-120; BF0072 and BN0019 from *Oryza*; IR0088 from *Echinochloa*), to limit the impact of population subdivision within lineages. The *Stenotaphrum-infecting*, lineage was not included in lineage-level analyses due to the small sample size.

Nucleotide diversity [84]. synonymous and non-synonymous nucleotide diversity, and population differentiation [45] were estimated using EGGLIB V3 [85] using classes COMPUTESTATS and CODINGDIVERSITY. Sites with more than 30% missing data were excluded. Orthogroups with less than 10 sequences (*nseff*<10, *nseff*being the average number of used samples among sites that passed the missing data filter) or shorter than 30bp (*lseff*<30, *lseff*being the number of sites used for analysis after filtering out sites with too many missing data) were excluded from computations. For analyses of polymorphism at secondary structure annotations, the cutoff on *Iseff*was set at 10bp.

For the computation of *d*_*N*_ / *d*_*S*_ and quantification of adaptive evolution, we used isolate NI919 of *Pyricularia* sp. LS [47, 48] as the outgroup (assembly GCA_004337975.1, European Nucleotide Archive). Genes were predicted in the outgroup assembly using Exonerate V2.2 Coding2genome [86]. For each gene, the query sequence was a *P. oryzae* sequence randomly selected among sequences with the fewest missing data. In parsing Exonerate output, we selected the sequence with the highest score, with a length greater than half the length of the query sequence.

The *d*_*N*_*/d*_*S*_ ratio was estimated using a maximum likelihood approach (runmode=-2, CodonFreq=2 in Codeml [87]), in pairwise comparisons of protein coding sequences (*i.e*., without using a phylogeny). For each *d*_*N*_*/d*_*S*_ we randomly selected 12 ingroup sequences and computed the average *d*_*N*_*/d*_*S*_ across the 12 ingroup/outgroup pairs.

Kruskal-Wallis tests were performed using the SCIPY.STATS.KRUSKAL library in PYTHON 3.7. Posthoc Mann-Whitney U-tests were performed using the SCIKIT_POSTHOCS library in PYTHON 3.7, with p-values adjusted using the Bonferroni-Holm method.

Amino acid change data was modeled using a binomial generalized linear mixed model with function GLMER with package LME4 version 1.1-32 in R version 4.1.2. Interactions between predictor variables were not significant and thus not included in the model presented in the Results section.

### Constructs for the transformation of fungal isolates

PCR products used for cloning were generated using the Phusion High-Fidelity DNA Polymerase (Thermo Fisher) and the primers listed in S7 Table. Details of the constructs are given in S8 Table. Briefly, the pSC642 plasmid (derived from the pCB1004 vector), containing a cassette for the expression of a gene of interest under the control of the *AVR-Pia* promoter (*pAVR-Pia*) and the *Neurospora crassa β-tubulin* terminator *(t-tub)*, was amplified by PCR with primers oML001 and oTK609 for the insertion of MAX genes listed in S9 Table. The MAX genes *Mo_US0071_000070 (MAX79), Mo_US0071_046730 (MAX89)* and *Mo_US0071_115900 ÇMAX83)*, amplified by PCR from genomic DNA of the *P. oryzae* isolate US0071, were inserted into this vector using the Gibson Assembly Cloning Kit (New England BioLabs). The final constructs were linearized using the KpnI restriction enzyme (Promega) before *P. oryzae* transformation.

### Plant and fungal growth conditions

Rice plants *(Oryza sativa)* were grown in a glasshouse in a substrate of 31% coconut peat, 30% Baltic blond peat, 15% Baltic black peat, 10% perlite, 9% volcanic sand, and 5% clay, supplemented with 3.5 g.L-^1^ of fertilizer (Basacote High K 6M, NPK 13-5-18). Plants were grown under a 12h-light photoperiod with a day-time temperature of 27□°C, night-time temperature of 21□°C, and 70% humidity. For spore production, the wild-type and transgenic isolates of *P. oryzae* Guy**11** were grown for 14 days at 25°C under a 12h-light photoperiod on rice flour agar medium (20 g.L^-1^ rice seed flour, 2.5 g.L^-1^ yeast extract, 1.5% agar, 500.000U penicillin g), supplemented with 240 μg.ml^-1^ hygromycin for transgenic isolates. For mycelium production, plugs of mycelium of *P. oryzae* Guy11 were grown in liquid medium (10 g.L^-1^ glucose, 3 g.L^-1^ KNO3, 2 g.L^-1^ KH2PO4, 2,5 g.L^-1^ yeast extract, 500 000U penicillin g) for 5 days at 25°C in the dark under agitation.

### Fungal transformation

Protoplasts from the isolate Guy11 of *P. oryzae* were transformed by heat shock with 10μg of KpnI-linearized plasmids for the expression of MAX effectors or RFP as described previously [88]. After two rounds of antibiotic selection and isolation of monospores, transformed isolates were genotyped by Phire Plant Direct PCR (Thermo Scientific) using primers described in S7 Table. The Guy11 transgenic isolates expressing *AVR-Pia* and *AVR1-CO39* were previously generated [50, 89].

### Fungal growth and infection assays

For the analysis of interaction phenotypes, leaves of three-week-old rice plants were spray-inoculated with conidial suspensions (40 000 conidia.ml^-1^ in water with 0.5% gelatin). Plants were incubated for 16 hours in the dark at 25°C and 95% relative humidity, and then grown for six days in regular growth conditions. Seven days after inoculation, the youngest leaf that was fully expanded at the time of inoculation was collected and scanned (Scanner Epson Perfection V370) for further symptoms analyses. Phenotypes were qualitatively classified according to lesion types: no lesion or small brown spots (resistance), small lesions with a pronounced brown border and a small gray center (partial resistance), and larger lesions with a large gray center or dried leaves (susceptibility). For the analysis of gene expression, plants were spray-inoculated with conidial suspensions at 50 000 conidia.ml^-1^ (in water with 0.5% gelatin), and leaves were collected three days after inoculation.

### RNA extraction and qRT-PCR analysis

Total RNA extraction from rice leaves or Guy11 mycelium and reverse transcription were performed as described by ref. [90]. Briefly, frozen leaves and mycelium were mechanically ground. RNA was extracted using TRI-reagent (Sigma-Aldrich) and chloroform separation. Denaturated RNA (5μg) was retrotranscribed and used for quantitative PCR using GoTaq qPCR Master Mix according to the manufacturer’s instructions (Promega) at a dilution of 1/10 for mycelium and 1/7 for rice leaves. The primers used are described in S7 Table. Amplification was performed as described by ref. [90] using a LightCycler480 instrument (Roche), and data were extracted using the instrument software. To calculate *MAX* gene expressions, the 2^-ΔΔCT^ method and primers measured efficiency were used. Gene expression levels are expressed relative to the expression of constitutive reference gene *MoEF1α*.

### Statistical analyses of phenotypic data

For expression comparison between Kitaake and Maratelli infection, all analyses were performed using R (www.r-proiect.org). The entire kinetic experiment was repeated three times with five biological replicates for each time point. For each variety, gene, and experimental replicate, values corresponding to the day post-inoculation with the highest median expression were extracted for statistical analyses. Expression data were not normally distributed so for each gene, differences between varieties were evaluated using non-parametric Mann-Whitney U-tests.

## SUPPORTING INFORMATION

S1 Table. Genomic assemblies with metadata.

S2 Table. Nomenclature of MAX effectors predicted in this study and in previous reports.

S3 Table. Presence/absence of MAX effector orthologs.

S4 Table. The expression of *MAX79, MAX83* and *MAX89* in Guy11 does not trigger recognition in a panel of rice varieties.

S5 Table. Gene average of summary statistics of polymorphism, differentiation and divergence.

S6 Table. π_*N*_ and *d*_*N*_/ *d*_*S*_ in different classes of secondary structure annotations for MAX effectors with π_*N*_*/*π_*S*_*>*1 and *d*_*N*_/ *d*_*S*_ *>1*., respectively.

S7 Table. Primers for cloning and expression analyses.

S8 Table. Vector constructs.

S9 Table. Sequences of the MAX effectors in the isolate US0071 that were used for the complementation of Guy11.

S1 Figure. Effect of assembly properties on the number of genes.

S2 Figure. Expression patterns of MAX effectors during rice infection.

S3 Figure. Differential expression levels of MAX effectors upon infection of two different rice cultivars.

S4 Figure. Nucleotide diversity (π), ratio of non-synonymous to synonymous nucleotide diversity (π_N_/π_S_), orthogroup frequency for MAX effectors, other secreted proteins, and other genes.

S5 Figure. Frequency of MAX effector orthogroups as a function of the frequency of the adjacent orthogroups in the genome.

S6 Figure. Analyses of population subdivision with sNMF.

S7 Figure. *MAX79, MAX83* and *MAX89* are expressed in the transgenic Guy11 isolates upon rice inoculation.

S8 Figure. *F*_*ST*_versus *π*_*N*_/*π*_*S*_ at MAX effectors

S9 Figure. Amino acid changes segregating in *P. oryzae* at MAX effectors with an avirulence function and MoToxB (first row), and MAX effectors with *π*_*N*_*/π*_*S*_*>2* (next rows); amino acid changes are shown in dark blue and known binding interfaces in light blue. Note that the *π*_*N*_/*π*_*S*_ ratio is >2 for the avirulence genes *AvrPiz-t* and AVR-Pik. Proteins are displayed twice, with one copy rotated 180 degrees around a vertical axis. The interface involved in binding with host proteins is known for AVR1-CO39, AVR-Pia, and AVR-Pik only ([50-53]).

S10 Figure. Secondary structure annotations of MAX effectors aligned with TM-ALIGN.

S1 Data. Summary statistics per orthogroup.

S2 Data. Summary statistics per MAX effector ortholog, species wide, and per lineage.

S3 Data. Structural properties and polymorphism of amino acids in MAX effectors.

S1 Text Fitting a generalized linear model to amino acid polymorphism data.

S2 Text Homology modeling procedure.

## REFERENCES

1. Schulze-Lefert P, Panstruga R. A molecular evolutionary concept connecting nonhost resistance, pathogen host range, and pathogen speciation. Trends in Plant Science. 2011;16(3):117–25. doi: 10.1016/j.tplants.2011.01.001.

2. Sánchez-Vallet A, Fouché S, Fudal I, Hartmann FE, Soyer JL, Tellier A, et al. The genome biology of effector gene evolution in filamentous plant pathogens. Annual review of phytopathology. 2018;56:21–40.

3. Haldane JBS. Disease and evolution. Ricerca Scient 1949;19:68–76.

4. Flor HH. Inheritance of pathogenicity in Melampsora lini. Phytopathology. 1942;32:653–69.

5. Barrett JA. Frequency-dependent selection in plant-fungal interactions. Philosophical Transactions of the Royal Society of London B, Biological Sciences. 1988;319(1196):473–83.

6. Bergelson J, Kreitman M, Stahl EA, Tian D. Evolutionary dynamics of plant R-genes. Science. 2001;292(5525):2281–5.

7. Brown JKM. Chance and selection in the evolution of barley mildew. Trends in Microbiology. 1994;2(12):470–5. PubMed PMID: 29.

8. Brown JKM. Durable resistance of crops to disease: a Darwinian perspective. Annual review of phytopathology. 2015;53:513–39.

9. Stahl EA, Dwyer G, Mauricio R, Kreitman M, Bergelson J. Dynamics of disease resistance polymorphism at the Rpm1 locus of Arabidopsis. Nature. 1999;400(6745):667–71.

10. Gladieux P, van Oosterhout C, Fairhead S, Jouet A, Ortiz D, Ravel S, et al. Extensive immune receptor repertoire diversity in disease-resistant rice landraces. bioRxiv. 2022:2022-12.

11. Bakker EG, Toomajian C, Kreitman M, Bergelson J. A genome-wide survey of R gene polymorphisms in Arabidopsis. The Plant cell. 2006;18(8):1803–18.

12. Bakker EG, Traw MB, Toomajian C, Kreitman M, Bergelson J. Low levels of polymorphism in genes that control the activation of defense response in Arabidopsis thaliana. Genetics. 2008;178(4):2031–43.

13. Ebert D, Fields PD. Host–parasite co-evolution and its genomic signature. Nature Reviews Genetics. 2020;21(12):754–68.

14. Ebbole DJ, Chen M, Zhong Z, Farmer N, Zheng W, Han Y, et al. Evolution and Regulation of a Large Effector Family of Pyricularia oryzae. Molecular Plant-Microbe Interactions. 2021;34(3):255–69.

15. Seong K, Krasileva KV. Computational structural genomics unravels common folds and novel families in the secretome of fungal phytopathogen Magnaporthe oryzae. Molecular Plant-Microbe Interactions. 2021;34(11):1267–80.

16. Seong K, Krasileva KV. Prediction of effector protein structures from fungal phytopathogens enables evolutionary analyses. Nature Microbiology. 2023;8(1):174–87.

17. de Guillen K, Ortiz-Vallejo D, Gracy J, Fournier E, Kroj T, Padilla A. Structure analysis uncovers a highly diverse but structurally conserved effector family in phytopathogenic fungi. PLoS Pathog. 2015;11(10):e1005228.

18. Savary S, Willocquet L, Pethybridge SJ, Esker P, McRoberts N, Nelson A. The global burden of pathogens and pests on major food crops. Nature ecology & evolution. 2019;3(3):430.

19. Fernandez J, Orth K. Rise of a cereal killer: the biology of Magnaporthe oryzae biotrophic growth. Trends in microbiology. 2018;26(7):582–97.

20. Couch BC, Fudal I, Lebrun M-H, Tharreau D, Valent B, van Kim P, et al. Origins of Host-Specific Populations of the Blast Pathogen Magnaporthe oryzae in Crop Domestication With Subsequent Expansion of Pandemic Clones on Rice and Weeds of Rice. Genetics. 2005;170(2):613–30. PubMed PMID: 61.

21. Pordel A, Ravel S, Charriat F, Gladieux P, Cros-Arteil S, Milazzo J, et al. Tracing the origin and evolutionary history of Pyricularia oryzae infecting maize and barnyard grass. Phytopathology. 2021;111(1):128–36.

22. Kato H, Yamamoto M, Yamaguchi-Ozaki T, Kadouchi H, Iwamoto Y, Nakayashiki H, et al. Pathogenicity, mating ability and DNA restriction fragment length polymorphisms of Pyricularia populations isolated from Gramineae, Bambusideae and Zingiberaceae plants. Journal of General Plant Pathology. 2000;66:30–47.

23. Urashima AS, Igarashi S, Kato H. Host range, mating type, and fertility of Pyricularia grisea from wheat in Brazil. Plant Disease. 1993;77(12):1211–6.

24. Igarashi S. Pyricularia em trigo. 1. Ocorrencia de Pyricularia sp noestado do Parana. Fitopatol Bras. 1986;11:351–2.

25. Milazzo J, Pordel A, Ravel S, Tharreau D. First scientific report of Pyricularia oryzae causing gray leaf spot disease on perennial ryegrass (Lolium perenne) in France. Plant Disease. 2019;103(5):1024-.

26. Islam MT, Croll D, Gladieux P, Soanes DM, Persoons A, Bhattacharjee P, et al. Emergence of wheat blast in Bangladesh was caused by a South American lineage of Magnaporthe oryzae. BMC biology. 2016;14(1):84.

27. Gladieux P, Condon B, Ravel S, Soanes D, Maciel JLN, Nhani A, et al. Gene Flow between Divergent Cereal- and Grass-Specific Lineages of the Rice Blast Fungus Magnaporthe oryzae. mBio. 2018;9(1).

28. Sweigard JA, Carroll AM, Kang S, Farrall L, Chumley FG, Valent B. Identification, Cloning, and Characterization of PWL2, a Gene for Host Species Specificity in the Rice Blast Fungus. The Plant cell. 1995;7(8):1221–33. PubMed PMID: 260.

29. Inoue Y, Vy TTP, Yoshida K, Asano H, Mitsuoka C, Asuke S, et al. Evolution of the wheat blast fungus through functional losses in a host specificity determinant. Science. 2017;357(6346):80–3.

30. Asuke S, Tanaka M, Hyon G-S, Inoue Y, Vy TTP, Niwamoto D, et al. Evolution of an Eleusine-specific subgroup of Pyricularia oryzae through a gain of an avirulence gene. Molecular Plant-Microbe Interactions. 2020;33(2):153–65.

31. Zheng Y, Zheng W, Lin F, Zhang Y, Yi Y, Wang B, et al. AVR1-CO39 is a predominant locus governing the broad avirulence of Magnaporthe oryzae 2539 on cultivated rice (Oryza sativa L.). Molecular plant-microbe interactions. 2011;24(1):13–7.

32. Simão FA, Waterhouse RM, Ioannidis P, Kriventseva EV, Zdobnov EM. BUSCO: assessing genome assembly and annotation completeness with single-copy orthologs. Bioinformatics. 2015;31(19):3210-2. 33.

33. Altschul SF, Madden TL, Schäffer AA, Zhang J, Zhang Z, Miller W, et al. Gapped BLAST and PSI-BLAST: a new generation of protein database search programs. Nucleic acids research. 1997;25(17):3389–402.

34. Finn RD, Clements J, Eddy SR. HMMER web server: interactive sequence similarity searching. Nucleic acids research. 2011;39(uppl_2):29–37.

35. Zhang Y, Skolnick J. TM-align: a protein structure alignment algorithm based on the TM-score. Nucleic acids research. 2005;33(7):2302–9.

36. Emms DM, Kelly S. OrthoFinder: solving fundamental biases in whole genome comparisons dramatically improves orthogroup inference accuracy. Genome biology. 2015;16(1):157.

37. Stamatakis A. RAxML version 8: a tool for phylogenetic analysis and post-analysis of large phylogenies. Bioinformatics. 2014;30(9):1312–3.

38. Yan X, Tang B, Ryder LS, MacLean D, Were VM, Eseola AB, et al. The transcriptional landscape of plant infection by the rice blast fungus Magnaporthe oryzae reveals distinct families of temporally co-regulated and structurally conserved effectors. The Plant cell. 2023;35(5):1360–85.

39. Fay JC, Wu C-I. Sequence divergence, functional constraint, and selection in protein evolution. Annual review of genomics and human genetics. 2003;4(1):213–35.

40. Frichot E, Mathieu F, Trouillon T, Bouchard G, François O. Fast and efficient estimation of individual ancestry coefficients. Genetics. 2014;196(4):973–83.

41. Huson DH, Bryant D. Application of Phylogenetic Networks in Evolutionary Studies. Molecular Biology and Evolution. 2006;23(2):254–67.

42. Tosa Y, Osue J, Eto Y, Oh H-S, Nakayashiki H, Mayama S, et al. Evolution of an avirulence gene, AVR1-CO39, concomitant with the evolution and differentiation of Magnaporthe oryzae. Molecular Plant-Microbe Interactions. 2005;18(11):1148–60.

43. Cesari S, Thilliez G, Ribot C, Chalvon V, Michel C, Jauneau A, et al. The rice resistance protein pair RGA4/RGA5 recognizes the Magnaporthe oryzae effectors AVR-Pia and AVR1-CO39 by direct binding. The Plant cell. 2013;25(4):1463-81. Epub 2013/04/04. doi: 10.1105/tpc.112.107201. PubMed PMID: 23548743; PubMed Central PMCID: PMCPMC3663280.

44. Okuyama Y, Kanzaki H, Abe A, Yoshida K, Tamiru M, Saitoh H, et al. A multifaceted genomics approach allows the isolation of the rice Pia-blast resistance gene consisting of two adjacent NBS-LRR protein genes. The Plant Journal. 2011;66(3):467–79.

45. Weir BS, Cockerham CC. Estimating F-statistics for the analysis of population structure. Evolution. 1984;38:1358–70. PubMed PMID: 283.

46. Hahn MW. Molecular population genetics: Oxford University Press; 2018.

47. Hirata K, Kusaba M, Chuma I, Osue J, Nakayashiki H, Mayama S, et al. Speciation in Pyricularia inferred from multilocus phylogenetic analysis. Mycological Research. 2007;111(7):799–808.

48. Gómez Luciano LB, Tsai IJ, Chuma I, Tosa Y, Chen Y-H, Li J-Y, et al. Blast fungal genomes show frequent chromosomal changes, gene gains and losses, and effector gene turnover. Molecular Biology and Evolution. 2019;36(6):1148–61.

49. Yang Z. Likelihood ratio tests for detecting positive selection and application to primate lysozyme evolution. Molecular biology and evolution. 1998;15(5):568–73.

50. Ortiz D, De Guillen K, Cesari S, Chalvon V, Gracy J, Padilla A, et al. Recognition of the Magnaporthe oryzae effector AVR-Pia by the decoy domain of the rice NLR immune receptor RGA5. The Plant cell. 2017;29(1):156–68.

51. Maqbool A, Saitoh H, Franceschetti M, Stevenson CEM, Uemura A, Kanzaki H, et al. Structural basis of pathogen recognition by an integrated HMA domain in a plant NLR immune receptor. Elife. 2015;4.

52. Guo L, Cesari S, de Guillen K, Chalvon V, Mammri L, Ma M, et al. Specific recognition of two MAX effectors by integrated HMA domains in plant immune receptors involves distinct binding surfaces. Proceedings of the National Academy of Sciences. 2018;115(45):11637–42.

53. Bentham AR, Petit-Houdenot Y, Win J, Chuma I, Terauchi R, Banfield MJ, et al. A single amino acid polymorphism in a conserved effector of the multihost blast fungus pathogen expands host-target binding spectrum. PLoS Pathogens. 2021;17(11):e1009957.

54. Clay K, Kover PX. The Red Queen hypothesis and plant/pathogen interactions. Annual review of Phytopathology. 1996;34(1):29–50.

55. Van Valen L. A new evolutionary law. 1973.

56. Yang Z, Bielawski JP. Statistical methods for detecting molecular adaptation. Trends in ecology & evolution. 2000;15(12):496–503.

57. Kourelis J, Marchal C, Posbeyikian A, Harant A, Kamoun S. NLR immune receptor–nanobody fusions confer plant disease resistance. Science. 2023;379(6635):934–9.

58. Chiapello H, Mallet L, Guerin C, Aguileta G, Amselem J, Kroj T, et al. Deciphering Genome Content and Evolutionary Relationships of Isolates from the Fungus Magnaporthe oryzae Attacking Different Host Plants. Genome Biol Evol. 2015;7(10):2896–912. Epub 2015/10/11. doi: 10.1093/gbe/evv187. PubMed PMID: 26454013; PubMed Central PMCID: PMCPMC4684704.

59. Soyer JL, El Ghalid M, Glaser N, Ollivier B, Linglin J, Grandaubert J, et al. Epigenetic control of effector gene expression in the plant pathogenic fungus Leptosphaeria maculans. PLoS genetics. 2014;10(3):e1004227.

60. Tang B, Yan X, Ryder LS, Cruz-Mireles N, Soanes DM, Molinari C, et al. Rgs1 is a regulator of effector gene expression during plant infection by the rice blast fungus Magnaporthe oryzae. bioRxiv. 2022:2022-09.

61. Fall LA, Salazar MM, Drnevich J, Holmes JR, Tseng M-C, Kolb FL, et al. Field pathogenomics of Fusarium head blight reveals pathogen transcriptome differences due to host resistance. Mycologia. 2019;111(4):563–73.

62. Sonah H, Zhang X, Deshmukh RK, Borhan MH, Fernando WGD, Belanger RR. Comparative transcriptomic analysis of virulence factors in Leptosphaeria maculans during compatible and incompatible interactions with canola. Frontiers in plant science. 2016;7:1784.

63. Arora S, Steed A, Goddard R, Gaurav K, O’Hara T, Schoen A, et al. A wheat kinase and immune receptor form host-specificity barriers against the blast fungus. Nature Plants. 2023:1–8.

64. Farman ML, Eto Y, Nakao T, Tosa Y, Nakayashiki H, Mayama S, et al. Analysis of the structure of the AVR1-Co39 avirulence locus in virulent rice-infecting isolates of Magnaporthe grisea. Molecular Plant-Microbe Interactions. 2002;15:6–16. PubMed PMID: 90.

65. Kang S, Sweigard Ja Fau - Valent B, Valent B. The PWL host specificity gene family in the blast fungus Magnaporthe grisea. Molecular Plant Microbe Interactions. 1995;(0894-0282 (Print)).

66. Brabham HJ, Gómez De La Cruz D, Were V, Shimizu M, Saitoh H, Hernández-Pinzón I, et al. Barley MLA3 recognizes the host-specificity determinant PWL2 from rice blast (M. oryzae). bioRxiv. 2022:2022-10.

67. Thierry M, Charriat F, Milazzo J, Adreit H, Ravel S, Cros-Arteil S, et al. Maintenance of divergent lineages of the Rice Blast Fungus Pyricularia oryzae through niche separation, loss of sex and post-mating genetic incompatibilities. PLoS pathogens. 2022;18(7):e1010687.

68. Martin M. Cutadapt removes adapter sequences from high-throughput sequencing reads. EMBnet journal. 2011;17(1):10–2.

69. Simpson JT, Wong K, Jackman SD, Schein JE, Jones SJM, Birol I. ABySS: a parallel assembler for short read sequence data. Genome research. 2009;19(6):1117–23.

70. Hoff KJ, Lange S, Lomsadze A, Borodovsky M, Stanke M. BRAKER1: unsupervised RNA-Seq-based genome annotation with GeneMark-ET and AUGUSTUS. Bioinformatics. 2015;32(5):767–9.

71. Abascal F, Zardoya R, Telford MJ. TranslatorX: multiple alignment of nucleotide sequences guided by amino acid translations. Nucleic acids research. 2010;38uppl_2):W7–W13.

72. Katoh K, Toh H. Recent developments in the MAFFT multiple sequence alignment program. Briefings in Bioinformatics. 2008;9(4):286–98. doi: 10.1093/bib/bbn013. PubMed PMID: ISI:000256756400004.

73. Castresana J. Selection of conserved blocks from multiple alignments for their use in phylogenetic analysis. Molecular biology and evolution. 2000;17(4):540–52.

74. Petersen TN, Brunak S, Von Heijne G, Nielsen H. SignalP 4.0: discriminating signal peptides from transmembrane regions. Nature methods. 2011;8(10):785–6.

75. Emanuelsson O, Nielsen H, Brunak S, Von Heijne G. Predicting subcellular localization of proteins based on their N-terminal amino acid sequence. Journal of molecular biology. 2000;300(4):1005–16.

76. Krogh A, Larsson B, Von Heijne G, Sonnhammer ELL. Predicting transmembrane protein topology with a hidden Markov model: application to complete genomes. Journal of molecular biology. 2001;305(3):567–80.

77. Robinson JT, Thorvaldsdóttir H, Turner D, Mesirov JP. igv. js: an embeddable JavaScript implementation of the Integrative Genomics Viewer (IGV). Bioinformatics. 2023;39(1):btac830.

78. Robinson JT, Thorvaldsdóttir H, Winckler W, Guttman M, Lander ES, Getz G, et al. Integrative genomics viewer. Nature biotechnology. 2011;29(1):24–6.

79. Webb B, Sali A. Protein Structure Modeling with MODELLER. Methods Mol Biol. 2021;2199(1940-6029 (Electronic)):239–55.

80. Zhou H, Zhou Y. Distance-scaled, finite ideal-gas reference state improves structure-derived potentials of mean force for structure selection and stability prediction. Protein science. 2002;11(11):2714–26.

81. Zhou H, Skolnick J. GOAP: a generalized orientation-dependent, all-atom statistical potential for protein structure prediction. Biophysical journal. 2011;101(8):2043–52.

82. Benkert P, Tosatto SCE, Schomburg D. QMEAN: A comprehensive scoring function for model quality assessment. Proteins: Structure, Function, and Bioinformatics. 2008;71(1):261–77.

83. Schrödinger L, DeLano W. PyMOL. 2020.

84. Tajima F. Evolutionary relationship of DNA sequences in finite populations. Genetics. 1983;105(2):437–60.

85. Siol M, Coudoux T, Ravel S, De Mita S. EggLib 3: A python package for population genetics and genomics. Molecular Ecology Resources. 2022;22(8):3176–87.

86. Slater GSC, Birney E. Automated generation of heuristics for biological sequence comparison. BMC bioinformatics. 2005;6(1):1–11.

87. Yang Z. PAML: a program package for phylogenetic analysis by maximum likelihood. Computer applications in the biosciences. 1997;13(5):555–6.

88. Villalba F, Collemare J, Landraud P, Lambou K, Brozek V, Cirer B, et al. Improved gene targeting in Magnaporthe grisea by inactivation of MgKU80 required for non-homologous end joining. Fungal Genetics and Biology. 2008;45(1):68–75.

89. Ribot C, Césari S, Abidi I, Chalvon V, Bournaud C, Vallet J, et al. The M agnaporthe oryzae effector AVR 1–CO 39 is translocated into rice cells independently of a fungal-derived machinery. The Plant Journal. 2013;74(1):1–12.

90. Pélissier R, Buendia L, Brousse A, Temple C, Ballini E, Fort F, et al. Plant neighbour-modulated susceptibility to pathogens in intraspecific mixtures. Journal of experimental botany. 2021;72(18):6570–80.

91. Frishman D, Argos P. Knowledge-based protein secondary structure assignment. Proteins: Structure, Function, and Bioinformatics. 1995;23(4):566–79.

