## Supplementary material for "Adaptive evolution in virulence effectors of the rice blast fungus *Pyricularia oryzae*": S1 Figure

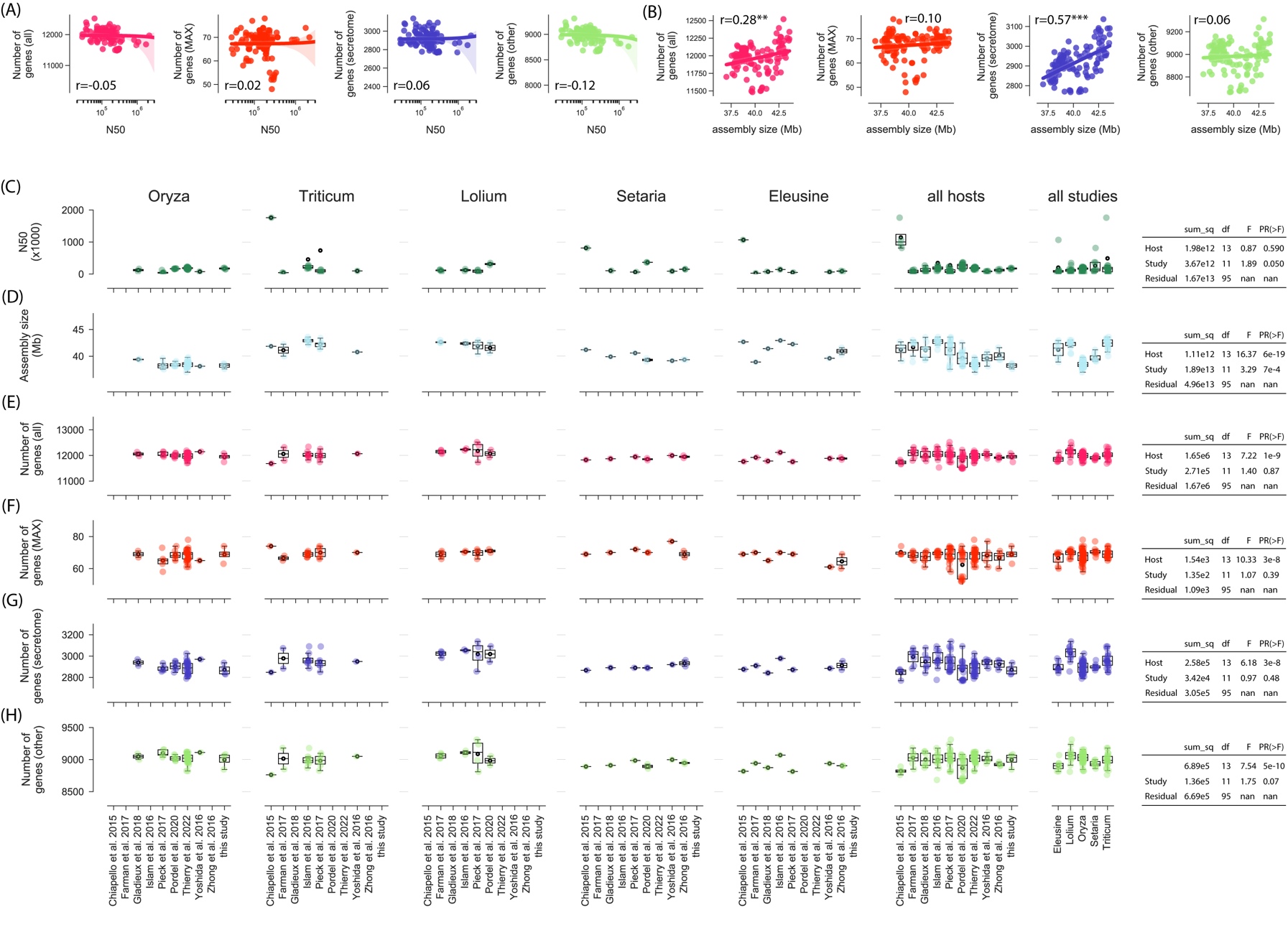

S1 Figure. Effect of assembly properties on the number of genes. (A) N50 versus number of genes encoding MAX effectors (‘MAX’), number of genes encoding secreted proteins (‘secretome’), number of other genes (‘other’), and total number of genes (‘all’); r is Pearson’s correlation coefficient (***p*<0.01; ****p*<0.001). (B) assembly size versus number of genes encoding MAX effectors (‘MAX’), number of genes encoding secreted proteins (‘secretome’), number of other genes (‘other’), and total number of genes (‘all’). (C) N50, assembly size, number of genes encoding MAX effectors (‘MAX’), number of genes encoding secreted proteins (‘secretome’), number of other genes (‘other’), and total number of genes (‘all’), as a function of the host of origin and study of origin of genomic data; tables report the results of type II analyses of variance (nan: not a number). In box plots, the black circle is the mean, the black line is the median. In box plots, the dashed black line is the mean, the solid black line is the median.
