## Supplementary material for "Adaptive evolution in virulence effectors of the rice blast fungus *Pyricularia oryzae*": S2 Figure

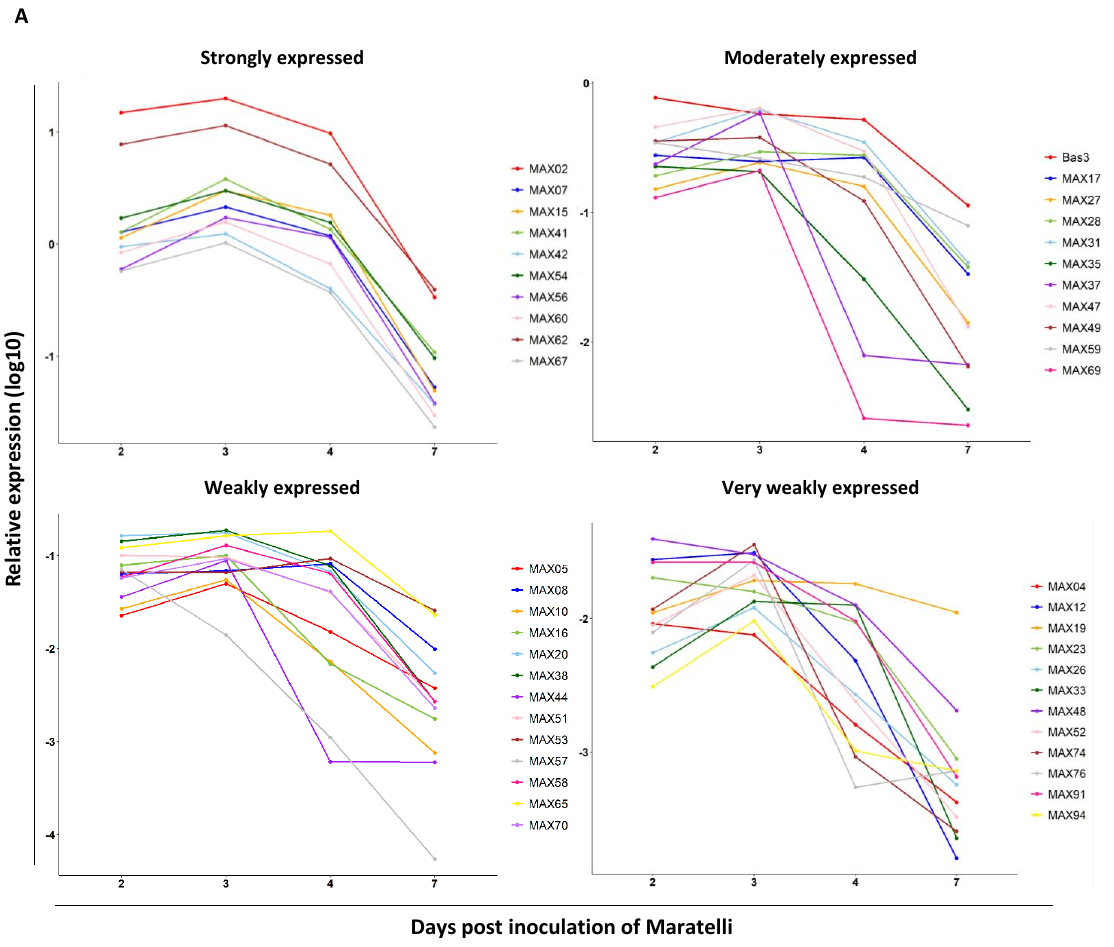

**S2 Figure. Expression patterns of MAX effectors during rice infection.**

Transcript levels of *MAX* effector genes, *Bas3* (biotrophy marker) and *MoEF1α* (Elongation Factor 1α, constitutive gene) were determined by qRT-PCR on leaves of *O. sativa* Maratelli **(A)** and Kitaake **(B)** infected with *P. oryzae* Guy11 2, 3, 4 and 7 days after inoculation. Values are given relative to the *MoEF1α* reference gene and were log2 transformed. Graph shows median values calculated from 3 independent experiments with 5 biological replicates each for very weakly (0,008-0,04), weakly (0,04-0,2), moderately (0.2-1) and strongly (>1) expressed MAX genes.
