## Supplementary material for "Adaptive evolution in virulence effectors of the rice blast fungus *Pyricularia oryzae*": S3 Figure

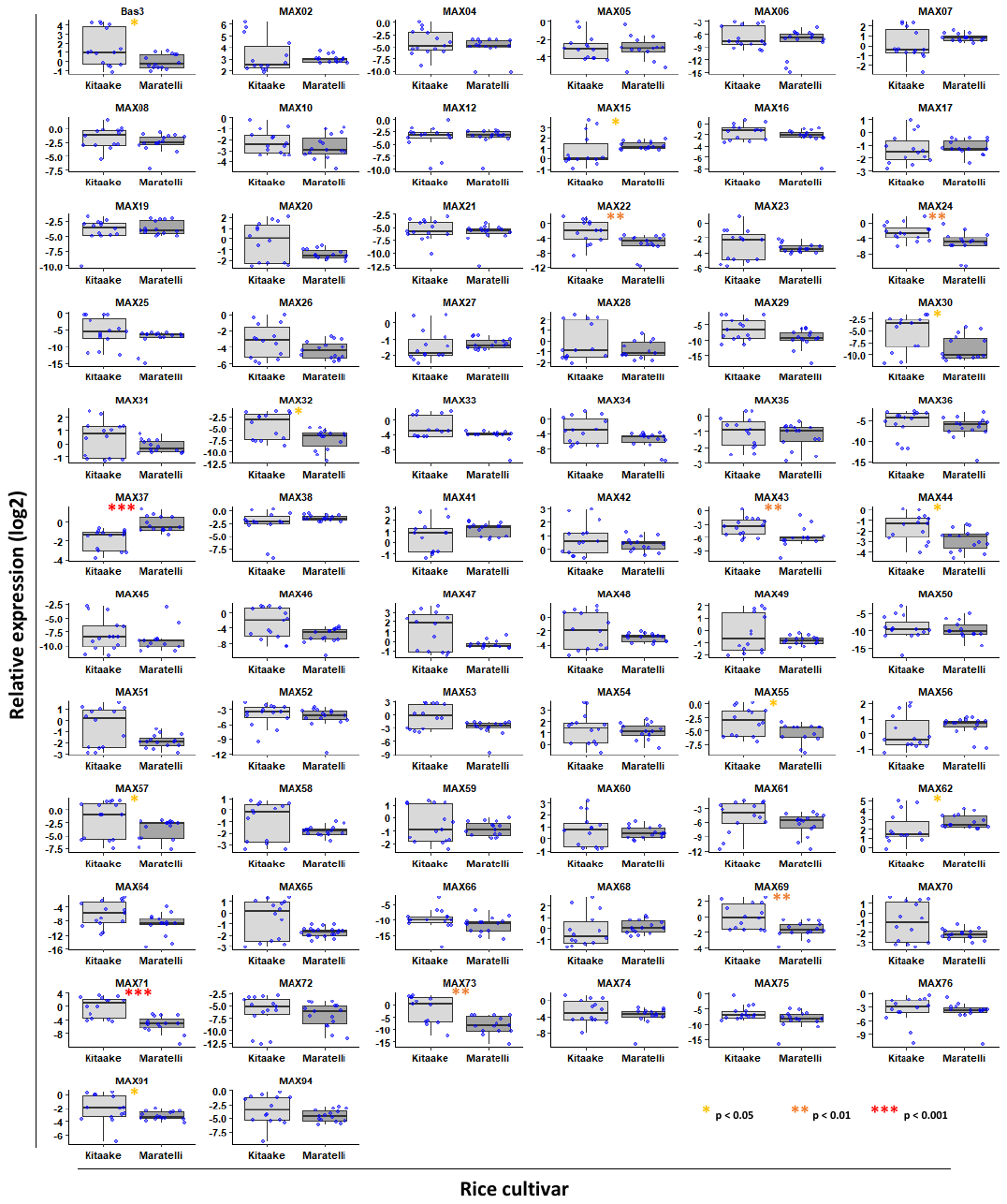

**S3 Figure. Differential expression levels of MAX effectors upon infection of two different rice cultivars**.

Transcript levels of *MAX* effectors were determined by qRT-PCR on leaves of *O. sativa* (Kitaake and Maratelli) infected with *P. oryzae* Guy11 2, 3, 4 and 7 days after inoculation. Values were calculated relative to the *MoEF1α* reference gene of *P. oryzae* and log2 transformed. For each variety, gene and replicate experiment, values corresponding to the day post inoculation with the highest median expression (calculated from 3-5 independent biological replicates) were kept. For each gene, boxplots show maximum expression values of all replicate experiments (blue dots) during Kitaake and Maratelli infection. Distribution was tested and Mann-Whitney U-tests were performed to assess for variability between the two conditions. Significance results are displayed with colored asterisks as follow: none (p >0.05), one yellow (p<0.05), two orange (p<0.01), three red (p<0.001).
