## Supplementary material for "Adaptive evolution in virulence effectors of the rice blast fungus *Pyricularia oryzae*": S4 Figure

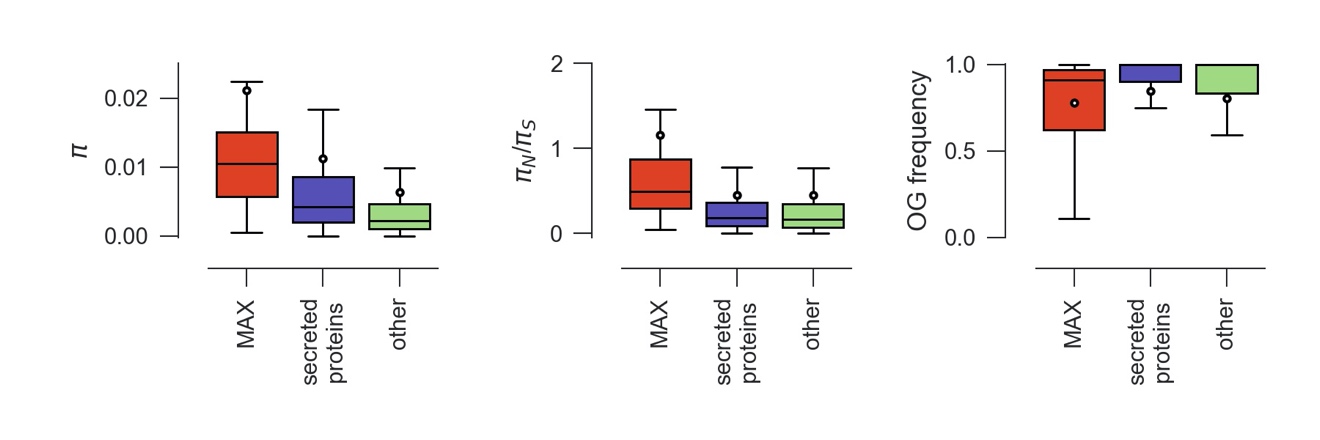

S4 Figure. Nucleotide diversity (π), ratio of non-synonymous to synonymous nucleotide diversity (π_N_/π_S_), orthogroup frequency for MAX effectors, other secreted proteins, and other genes. All differences were statistically significant at *p*<0.05 (Kruskal-Wallis tests post-hoc Mann-Whitney U-tests). In box plots, the circle is the mean, the solid black line is the median.
