## Supplementary material for "Adaptive evolution in virulence effectors of the rice blast fungus *Pyricularia oryzae*": S5 Figure

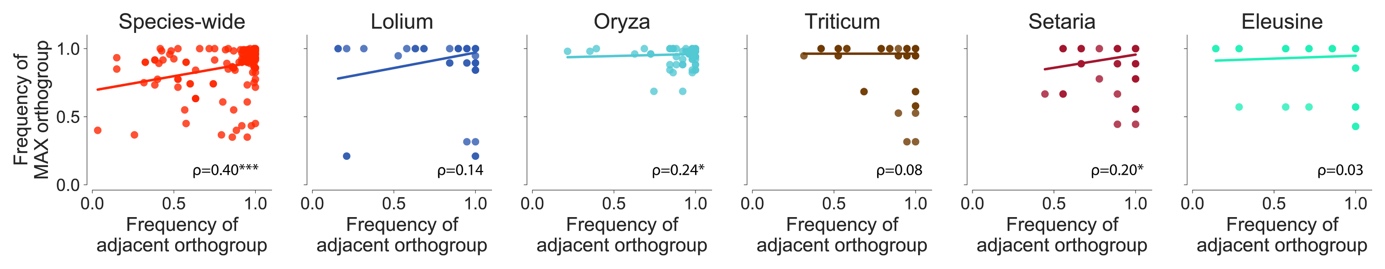

S5 Figure. Frequency of MAX effector orthogroups as a function of the frequency of the adjacent orthogroups in the genome. ρ is Spearman’s rank-order correlation statistic. ****p*<0.001; **p*<0.05.
