## Supplementary material for "Adaptive evolution in virulence effectors of the rice blast fungus *Pyricularia oryzae*": S6 Figure

(A)

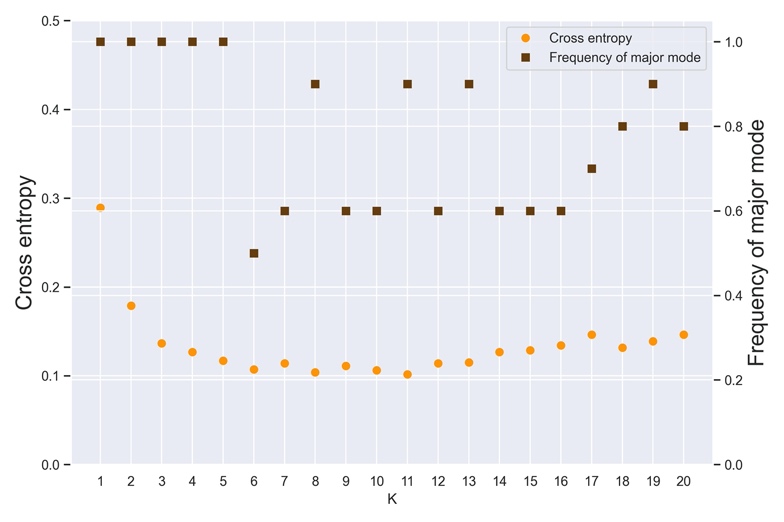

(B)

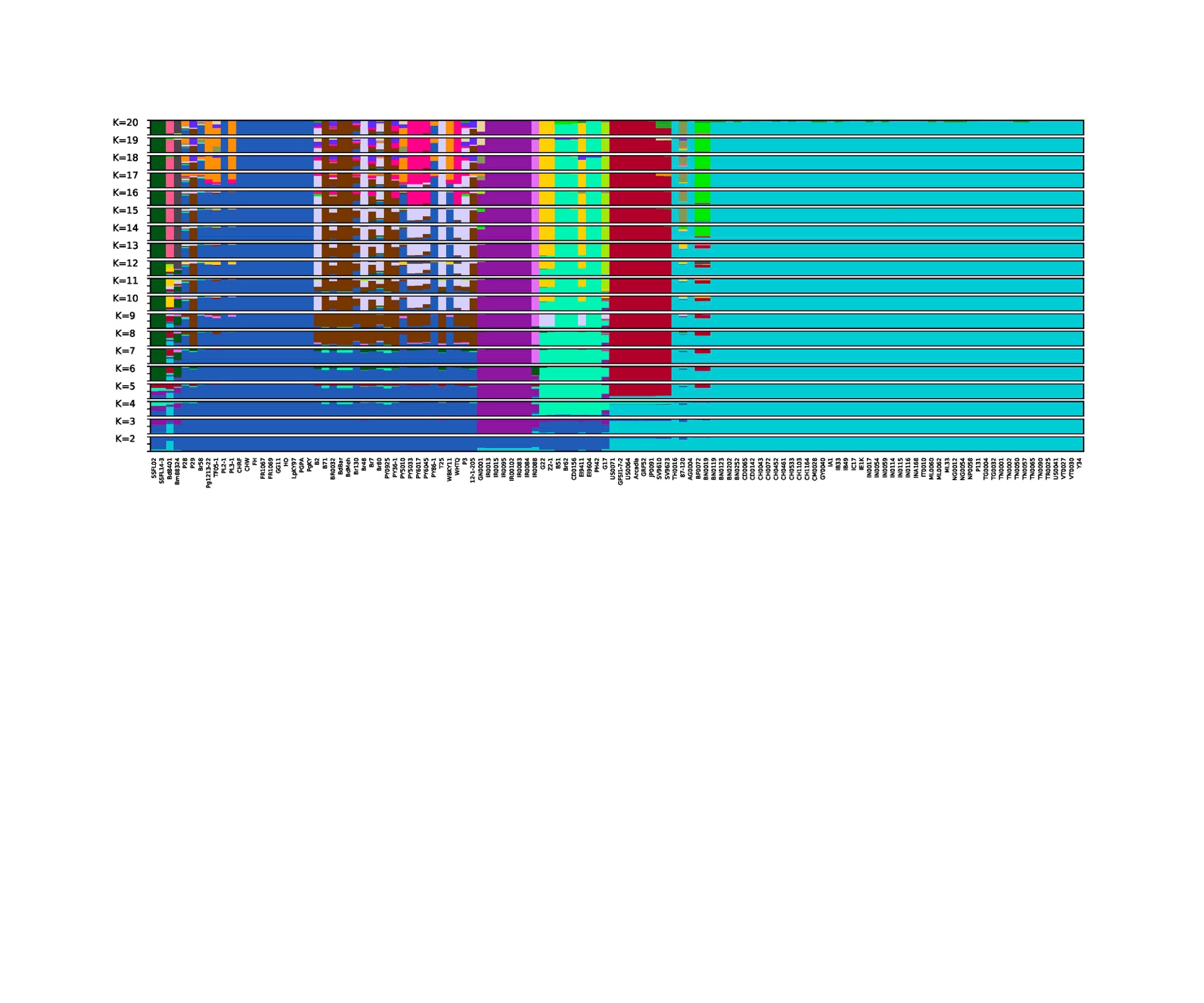

S6 Figure. Analyses of population subdivision with sNMF. (A) Cross-entropy and frequency of the major mode as a function of the number of clusters K in sNMF analyses of population subdivision (B) Genetic ancestry proportions for individual isolates in K=2 to K=20 ancestral populations. Each isolate is depicted as a horizontal bar divided into K segments representing the proportion of ancestry in K inferred ancestral populations. Genetic ancestry proportions were inferred from 6780 polymorphisms at four-fold degenerate synonymous sites identified in coding sequences of single-copy core orthologs (one polymorphism without missing data randomly chosen per ortholog). The major mode refers to the clustering solution with the highest frequency among 10 replicate runs of the sNMF algorithm for each K. Only major modes are represented in (B).
