## Supplementary material for "Adaptive evolution in virulence effectors of the rice blast fungus *Pyricularia oryzae*": S7 Figure

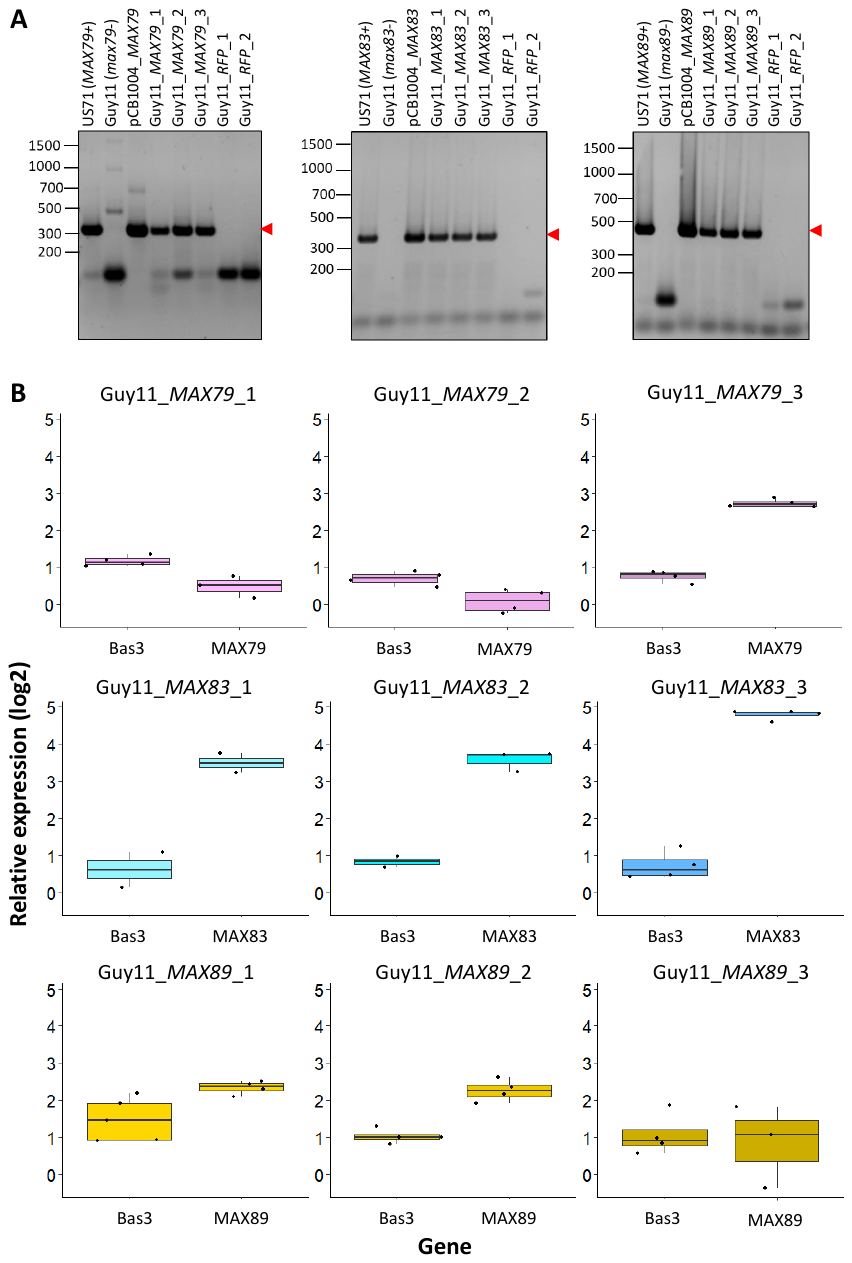

**S7 Figure. *MAX79, MAX83* and *MAX89* are expressed in the transgenic Guy11 isolates upon rice inoculation.**

**(A)** Genotyping by PCR of transgenic *P. oryzae* Guy11 complemented with *MAX79*, *MAX83* or *MAX89*. Expected PCR product sizes are 341bp (*MAX79*), 383bp (*MAX83*) and 454bp (*MAX89*). Red arrows show expected PCR products. The US0071 isolate (carrying *MAX79*, *MAX83* and *MAX89*) and pCB1004 plasmids (used for transformation) are used as positive controls. Wild type Guy11 isolate (lacking *MAX79*, *MAX83* and *MAX89*) and Guy11 complemented with *RFP* are used as negative controls. **(B)** Box plots showing the quantification of foreign *MAX* expression in transgenic Guy11 isolates. Transcript levels of *MAX79*, *MAX83*, *MAX89*, *Bas3* (Biotrophy marker) and *MoEf1α* (Elongation Factor 1α) were determined by qRT-PCR on 2-5 independent leaves of *O. sativa* (Maratelli) infected with transgenic *P. oryzae* Guy11 transformed with *MAX79*, *MAX83* or *MAX89* three days after inoculation. *Bas3* is considered strongly expressed during biotrophy and is used here as a mean of comparison for the expression level of transgenes. Values are given relative to the *MoEf1α* reference gene and were log2 transformed.
