## Supplementary material for "Adaptive evolution in virulence effectors of the rice blast fungus *Pyricularia oryzae*": S8 Figure

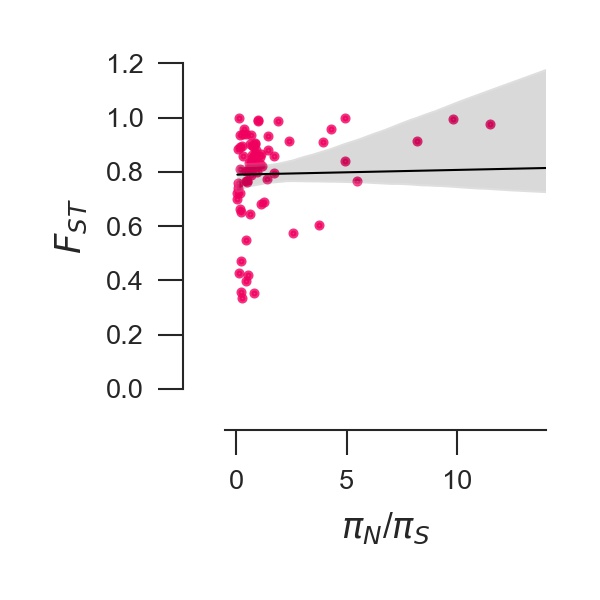

S8 Figure. *F_ST_* versus *π_N_/π_S_* at MAX effectors, with regression model y ~ x and 95% confidence interval for that regression (as estimated using regplot function in Seaborn package with Python3.7).
