## Supplementary material for "Adaptive evolution in virulence effectors of the rice blast fungus *Pyricularia oryzae*": S9 Figure

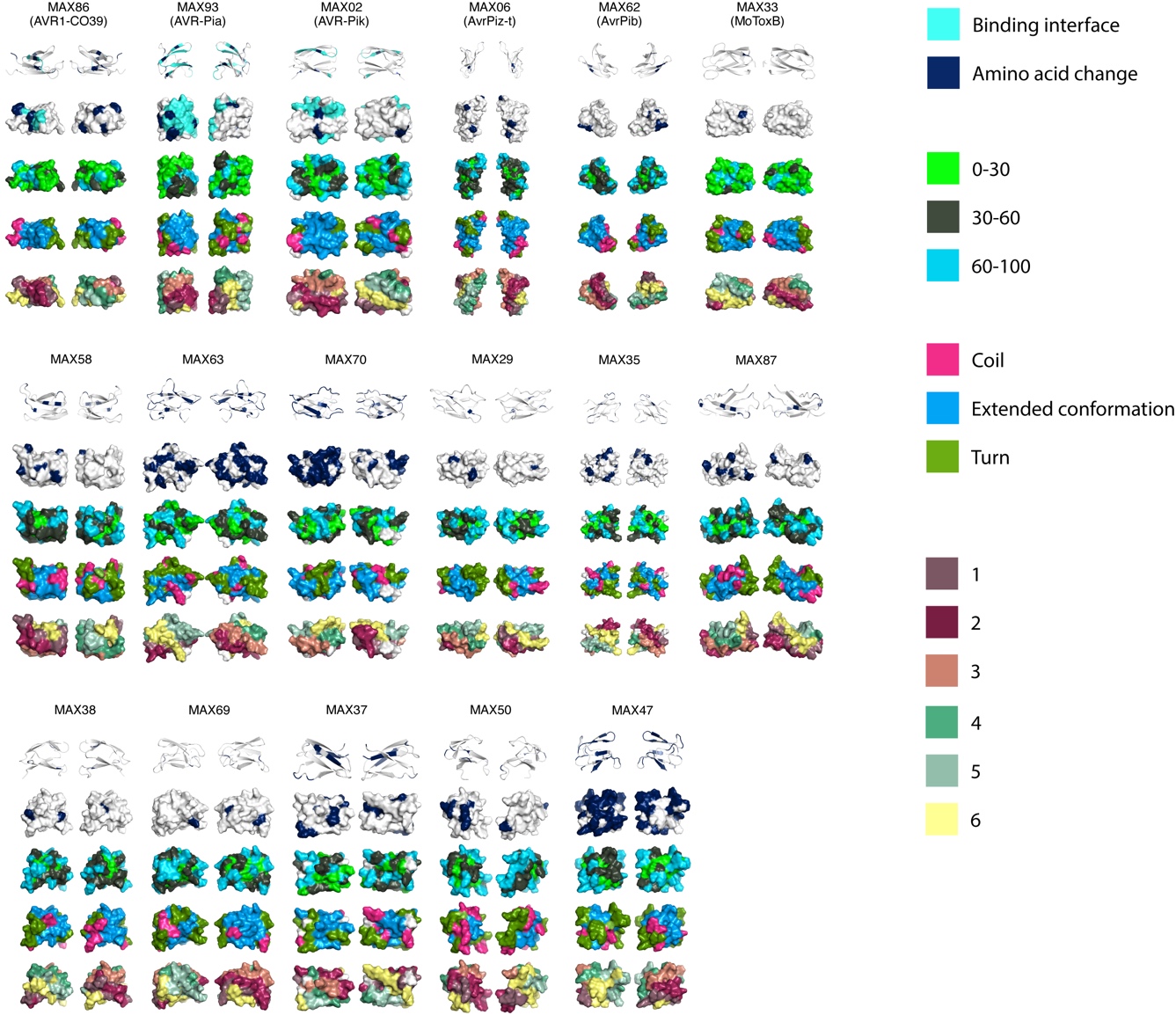

S9 Figure. Amino acid changes segregating in *P. oryzae* at MAX effectors with an avirulence function and MoToxB (first row), and MAX effectors with *π_N_/π_S_*>2 (next rows); amino acid changes are shown in dark blue and known binding interfaces in light blue. Note that the *π_N_/π_S_* ratio is >2 for the avirulence genes *AvrPiz-t* and AVR-Pik. Proteins are displayed twice, with one copy rotated 180 degrees around a vertical axis. The interface involved in binding with host proteins is known for AVR1-CO39, AVR-Pia, and AVR-Pik only ([54-57]).
