## Supplementary material for "Adaptive evolution in virulence effectors of the rice blast fungus *Pyricularia oryzae*": S1 Text

### S1 Text. Fitting a generalized linear model to amino acid polymorphism data.

We modeled the probability that an amino acid is polymorphic.

The data used is in S3 Data.txt.

We use the glmer function in R 4.1.2. to fit a binomial regression to our data.

```
library(lme4)
library(boot)
mydata = read.table("input.txt",header=T) # =S3 Data.txt without its first line
mydata$domain <- as.factor(as.vector(mydata$domain))
mydata$accessibility <- as.factor(as.vector(mydata$accessibility))
mydata$feature <- as.factor(as.vector(mydata$feature))
mymodel<-glmer(polymorphic~domain + feature + accessibility + (1|MAX), family=binomial, data=mydata)
summary(mymodel)
```

```
Generalized linear mixed model fit by maximum likelihood (Laplace Approximation) ['glmerMod']
 Family: binomial  ( logit )
Formula: polymorphic ~ domain + feature + accessibility + (1 | MAX)
   Data: mydata

     AIC      BIC   logLik deviance df.resid
  3469.2   3542.7  -1723.6   3447.2     5899

Scaled residuals:
    Min      1Q  Median      3Q     Max
-1.3210 -0.3352 -0.2478 -0.1719  7.3324

Random effects:
 Groups Name        Variance Std.Dev.
 MAX    (Intercept) 1.358    1.165
Number of obs: 5910, groups:  MAX, 91

Fixed effects:
                    Estimate Std. Error z value Pr(>|z|)
(Intercept)         -2.90333    0.21013 -13.817  < 2e-16 ***
domain2              0.07997    0.15707   0.509   0.6106
domain3              0.19218    0.16091   1.194   0.2324
domain4              0.04527    0.16653   0.272   0.7857
domain5              0.09878    0.16107   0.613   0.5397
domain6             -0.03777    0.15813  -0.239   0.8112
featureE             0.03171    0.13590   0.233   0.8155
featureT            -0.04377    0.13134  -0.333   0.7390
accessibility30-60   0.29119    0.12006   2.425   0.0153 *
accessibility60-100  0.58613    0.12602   4.651  3.3e-06 ***
---
Signif. codes:  0 ‘***’ 0.001 ‘**’ 0.01 ‘*’ 0.05 ‘.’ 0.1 ‘ ’ 1

Correlation of Fixed Effects:
            (Intr) doman2 doman3 doman4 doman5 doman6 featrE featrT a30-60
domain2     -0.307
domain3     -0.292  0.525
domain4     -0.311  0.499  0.486
domain5     -0.359  0.513  0.503  0.473
domain6     -0.373  0.527  0.513  0.484  0.520
featureE    -0.514 -0.078 -0.089 -0.079  0.059  0.071
featureT    -0.391 -0.043 -0.069 -0.039 -0.035  0.086  0.638
accssb30-60 -0.291 -0.174 -0.119 -0.067 -0.093 -0.134  0.141  0.006
accss60-100 -0.366 -0.105 -0.131 -0.053 -0.070 -0.133  0.307 -0.014  0.573
```

Then, we can estimate prediction intervals for the three subclasses of the factor "accessibility" using the boot package:

```
new.df <- mydata[sample(nrow(mydata), replace = TRUE), ]
new.df$accessibility <- as.factor("0-30")
my.bootstrap.predictions.f <- function(data, indices){
  return(mean(predict(mymodel, newdata = data[indices, ], type = "response", allow.new.levels=TRUE),
na.rm=TRUE))
}
my.boot.obj <- boot(data = new.df[sample(nrow(new.df), 20000, replace = TRUE), ], statistic =
my.bootstrap.predictions.f, R = 2)
quantile(my.boot.obj[[2]], c(0.025, 0.975))
```

```
      2.5%      97.5%
0.08314063 0.08459802
```

```
new.df <- mydata[sample(nrow(mydata), replace = TRUE), ]
new.df$accessibility <- as.factor("30-60")
my.bootstrap.predictions.f <- function(data, indices){
  return(mean(predict(mymodel, newdata = data[indices, ], type = "response", allow.new.levels=TRUE),
na.rm=TRUE))
}
my.boot.obj <- boot(data = new.df[sample(nrow(new.df), 20000, replace = TRUE), ], statistic =
my.bootstrap.predictions.f, R = 2)
quantile(my.boot.obj[[2]], c(0.025, 0.975))
```

```
     2.5%     97.5%
0.1057203 0.1065073
```

```
new.df <- mydata[sample(nrow(mydata), replace = TRUE), ]
new.df$accessibility <- as.factor("60-100")
my.bootstrap.predictions.f <- function(data, indices){
  return(mean(predict(mymodel, newdata = data[indices, ], type = "response", allow.new.levels=TRUE),
na.rm=TRUE))
}
my.boot.obj <- boot(data = new.df[sample(nrow(new.df), 20000, replace = TRUE), ], statistic =
my.bootstrap.predictions.f, R = 2)
quantile(my.boot.obj[[2]], c(0.025, 0.975))
```

```
     2.5%     97.5%
0.1308136 0.1311139
```

We can also compute the observed rates of amino acid polymorphism:

```
newdata <- mydata[ which(mydata$accessibility=="0-30"), ]
mean(newdata$polymorphic)
```

```
[1] 0.08674464
```

```
newdata <- mydata[ which(mydata$accessibility=="30-60"), ]
mean(newdata$polymorphic)
```

```
[1] 0.1052364
```

```
newdata <- mydata[ which(mydata$accessibility=="60-100"), ]
mean(newdata$polymorphic)
```

```
[1] 0.1295611
```
