## Supplementary material for "Adaptive evolution in virulence effectors of the rice blast fungus *Pyricularia oryzae*": S2 Text

S2 Text. Homology modeling procedure (multiple template-driven modeling and TM-pred score)

*Modeled sequences and 3D templates:*

Eight experimental structures with MAX-like folds were selected as 3D templates for homology modeling (PDB identifiers of the templates: 6R5J, 2MM0, 2MM2, 2MYW, 2LW6, 5A6W, 5Z1V, 5ZNG). They were aligned using the structural superposition program TM-align [1].A multiple sequence alignment **A** and a phylogenetic tree **T** of all MAX effectors and 3D templates were built using MAFFT v7 [2] and FastME v2 [3] software, respectively.

For each of the MAX orthologous groups, one representative protein was selected and homology models of this 1D query relatively to each 3D template were built using Modeller [5] with many alternative query-template threading alignments.

*Query-template threading.*

Alternative alignments between each 1D query and each 3D template were generated in the following way:

i) A 1D profile was built by selecting the ***N*** closest homologs to the 1D query in the global alignment ***A*** according to the phylogenetic tree ***T***. A 3D profile was obtained similarly by aligning the ***N*** closest template homologs.

ii) Optimal alignment scores E_N_(*i,j*) from the N-ter positions to positions (*i,j*) of the 1D and 3D profiles were built using the following dynamic programming equation :

E_N_(i,j) = max { E_N_(*i-1,j-1*)+E_1D_(*i,j*)+E_3D_(*i,j*), E_N_(*i-1,j*)+E_2D_(*j*), E_N_(*i,j-1*)+E_2D_(*j*) }	[1]

E_1D_(*i,j*) is the BLOSUM62 score [4] averaged over all matching residue pairs from the query and template profile positions *i* and *j*. E_3D_(*i,j*) measures the average compatibility between 1D profile residues at position *i* and template secondary structure and solvent accessibility at position *j* according to a statistical potential derived from a representative set of PDB structures. E_2D_(*j*) is a negative gap cost which penalizes alignment indels depending on the secondary structure at template position *j* (higher penalties are set in the secondary structure core shared by all templates).

iii) Optimal alignment scores E_C_(*i,j*) from the C-ter positions were obtained by dynamic programming (eq. 1) as in step ii) but in reverse sequence direction.

iv) The optimal alignment scoring matrices ***E***_N_ and ***E***_C_ were added, resulting in a matrix ***E*** whose cell (*i,j*) is measuring the score of the best alignment passing through the residue *i* of the 1D query and the residue *j* of the 3D template.

v) The query-template alignment with highest score in ***E*** is saved and its path is masked in the matrix ***E***. This step is repeated to generate a set of alternative alignments between the 1D query and the 3D template.

vi) Two alternative query-template alignments are randomly chosen. If they share pairs of aligned residues, one of these pairs is randomly chosen, namely residue *i* of the 1D query and residue *j* of the 3D template. Then the two alternative alignments are hybridized around the pivot (*i,j*) by merging the N-ter path (1..*i*)/(1..*j*) of the first alignment and the C-ter path (*i..l*_Q_)/(*j..l*_T_) of the second alignment, where *l*_Q_ is the query length and *l*_T_ is the template length (Figure 1). If the resulting alignment is new, it replaces the alignment with lowest score in the hybridization pool. This hybridization step is repeated to progressively improve the average score of the alternative alignments.

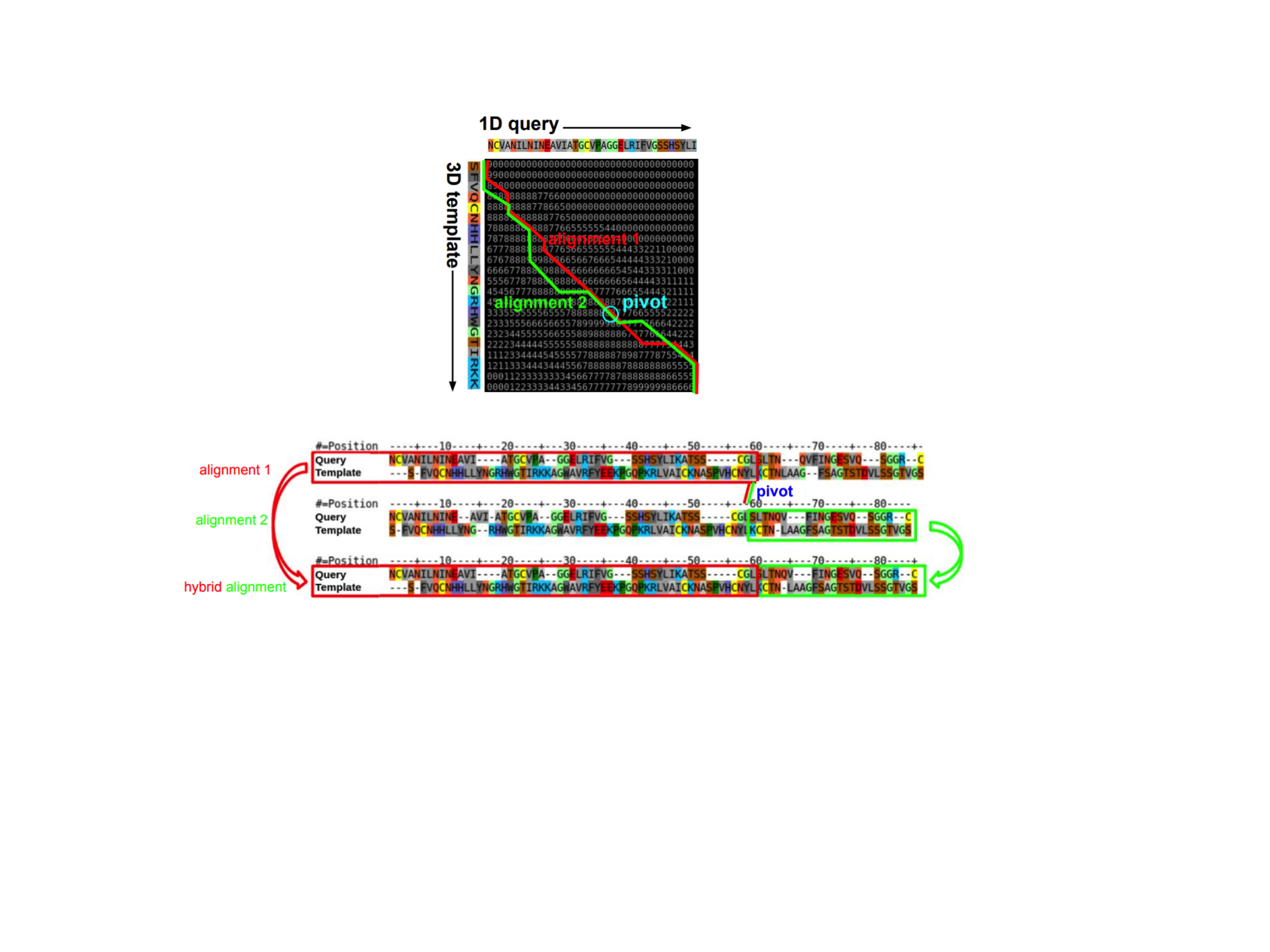

***Figure 1.*** *Hybridization of two alignments shown by red and green boxes, respectively, around a shared pivot position highlighted in blue.*

*Model construction.* Using the above procedure, 20 alternative alignments were generated for each pair of query and 3D template sequences, respectively, each of them was aligned with their closest homologs. The number ***N*** of homologs aligned in each profile was successively set to 1, 2, 4, 8, 16, 32, 64, and 128. Since eight templates were used, a maximum of 20x8x8 = 1280 alternative alignments were generated for each 1D query. The actual number of non-redundant alignments was actually lower since profiles with different numbers of homologs could generate the same query-template alignment. For each query-template alternative alignment, a structural model of the query was generated using MODELLER [5].

*TM-pred scoring.* We used this approach to build a learning set of 3D models by comparing all pairs of the eight templates with known structures. Each template was first discarded from the pool of 3D templates and introduced as a query sequence. For each query-template pair, sub-optimal alignments and their corresponding homology models were constructed as explained above. The accuracy of each query homology model was evaluated using 6 different scores (DFIRE [6], GOAP [7], QMEAN [8], E_1D_, E_2D_, E_3D_). E_1D_, E_2D_, E_3D_ are the cumulated scores from the dynamic programming equation [1] along the query-template alignment path. The weights of a linear combination of the 6 scores were tuned by linear regression to optimally fit the TM-score observed when superposing by TM-ALIGN [1] the query homology models with their corresponding experimental structures, which varies from 0 (distant) to 1 (r.m.s.d of 0.0). We termed this optimized score TM-pred which is a prediction of the TM-score which could be greater than 1. Higher values indicated better models. Values near 1 indicated quasi-perfect models. TM-pred was used to rank the models derived by MODELLER from the sub-optimal alignments built for each OG representative sequence. For each MAX orthologous group, the five structural models with highest TM-pred scores were collected and displayed here: <https://pat.cbs.cnrs.fr/magmax/model/>.

**REFERENCES**

1.	Zhang Y, Skolnick J. TM-align: a protein structure alignment algorithm based on the TM-score. Nucleic Acids Res. 2005;33: 2302–2309. doi:10.1093/nar/gki524

2.	Katoh K, Standley DM. MAFFT Multiple Sequence Alignment Software Version 7: Improvements in Performance and Usability. Mol Biol Evol. 2013;30: 772–780. doi:10.1093/molbev/mst010

3.	Lefort V, Desper R, Gascuel O. FastME 2.0: A Comprehensive, Accurate, and Fast Distance-Based Phylogeny Inference Program: Table 1. Mol Biol Evol. 2015;32: 2798–2800. doi:10.1093/molbev/msv150

4.	Henikoff S, Henikoff JG. Amino acid substitution matrices from protein blocks. Proc Natl Acad Sci. 1992;89: 10915–10919. doi:10.1073/pnas.89.22.10915

5.	Webb B, Sali A. Protein Structure Modeling with MODELLER. Methods Mol Biol. 2020;2199: 239–255. doi:10.1007/978-1-0716-0892-0_14

6.	Zhou H, Zhou Y. Distance-scaled, finite ideal-gas reference state improves structure-derived potentials of mean force for structure selection and stability prediction. Protein Sci. 2009;11: 2714–2726. doi:10.1110/ps.0217002

7.	Zhou H, Skolnick J. GOAP: A Generalized Orientation-Dependent, All-Atom Statistical Potential for Protein Structure Prediction. Biophys J. 2011;101: 2043–2052. doi:10.1016/j.bpj.2011.09.012

8.	Benkert P, Tosatto SCE, Schomburg D. QMEAN: A comprehensive scoring function for model quality assessment. Proteins Struct Funct Bioinforma. 2008;71: 261–277. doi:10.1002/prot.21715
