## Supplementary material for "Adaptive evolution in virulence effectors of the rice blast fungus *Pyricularia oryzae*": S5 Table

S5 Table. Gene average of summary statistics of polymorphism, differentiation and divergence.

| Dataset | Gene set | *π* | *F_ST_* | *π_N_* | *π_S_* | *π_N_/π_S_* | *d_N_/d_S_* |
| --- | --- | --- | --- | --- | --- | --- | --- |
| Species-wide | MAX | 0.0104 | 0.786 | 0.0078 | 0.0102 | 1.826 | 0.977 |
|  | Secretome | 0.0079 | 0.732^a^ | 0.0028 | 0.0117 | 0.461 | 0.711 |
|  | Other | 0.0049 | 0.733^a^ | 0.0017 | 0.0081 | 0.461 | 0.584 |
| Oryza lineage | MAX | 0.0015 | NA | 0.0012 | 0.0025 | 1.011 | 0.516 |
|  | Secretome | 0.0018 | NA | 0.0006 | 0.0022 | 0.438^a^ | 0.270 |
|  | Other | 0.0013 | NA | 0.0004 | 0.0016 | 0.493^a^ | 0.261 |
| Lolium lineage | MAX | 0.0015^ab^ | NA | 0.0037 | 0.0057^a^ | 0.863 | 0.521 |
|  | Secretome | 0.0024^a^ | NA | 0.0016 | 0.0061^a^ | 0.339 | 0.282 |
|  | Other | 0.0013^b^ | NA | 0.0010 | 0.0045 | 0.289 | 0.230 |
| Eleusine lineage | MAX | 0.0069^a^ | NA | 0.0065 | 0.0158^a^ | 0.872 | 0.539 |
|  | Secretome | 0.0044^a^ | NA | 0.0027 | 0.0103^a^ | 0.306^a^ | 0.258 |
|  | Other | 0.0025 | NA | 0.0016 | 0.0073 | 0.303^a^ | 0.223 |
| Setaria lineage | MAX | 0.0041^ab^ | NA | 0.0043 | 0.0184 | 0.384^a^ | 0.541 |
|  | Secretome | 0.0018^a^ | NA | 0.0012 | 0.0048 | 0.378^a^ | 0.254 |
|  | Other | 0.0011^b^ | NA | 0.0008 | 0.0032 | 0.385^a^ | 0.229 |
| Wheat lineage | MAX | 0.0034^a^ | NA | 0.0043 | 0.0103^a^ | 0.674 | 0.512 |
|  | Secretome | 0.0039^a^ | NA | 0.0019 | 0.0081^a^ | 0.308^a^ | 0.270 |
|  | Other | 0.0021 | NA | 0.0013 | 0.0059 | 0.289^a^ | 0.238 |

*π* (nucleotide diversity per bp), *F_ST_* (Weir and Cockerham’s estimator of population differentiation: https://doi.org/10.2307/2408641), π_S_ (synonymous nucleotide diversity per bp), π_N_ (non-synonymous nucleotide diversity per bp), π_N_/π_S_ (the ratio of non-synonymous to synonymous nucleotide diversity), d_N_/d_S_ (the ratio of non-synonymous to synonymous rates of substitutions). Shared superscripts indicate non-significant differences (post-hoc Mann-Whitney tests with Bonferroni-Holm correction, p-value>0.05). NA, non-applicable, because *F_ST_* is only calculated between lineages.
