## Supplementary material for "Adaptive evolution in virulence effectors of the rice blast fungus *Pyricularia oryzae*": S6 Table

S6 Table. *π_N_* and *d_N_/d_S_* in different classes of secondary structure annotations for MAX effectors with *π_N_/π_S_*>1 and *d_N_/d_S_*>1, respectively. NA, non-applicable.

| Statistic | MAX effector | Structural features | | | Solvent accessibility classes | | | Structural domains | | | | | |
| --- | --- | --- | --- | --- | --- | --- | --- | --- | --- | --- | --- | --- | --- |
|  |  | C | E | T | 0-30% | 30-60% | 60-100% | 1 | 2 | 3 | 4 | 5 | 6 |
| *π_N_* | OG0000073_2 | NA | 0.0059 | NA | 0.00485 | 0.00799 | NA | NA | 0.01135 | 0.01217 | NA | NA | NA |
|  | OG0000167_1 | NA | 0.00771 | 0.01415 | NA | 0.01181 | 0.01383 | NA | 0.02946 | NA | NA | NA | 0.01993 |
|  | OG0008177 | 0.01224 | 0.0031 | 0.02273 | 0.00393 | 0.02874 | 0.0121 | 0.00766 | 0.01357 | 0.02224 | NA | 0.02423 | 0.01767 |
|  | OG0008883 | 0.00198 | 0.00623 | 0.01466 | 0.00407 | 0.00222 | 0.02394 | 0.00413 | 0.01013 | 0.00563 | 0.02187 | 0.00659 | 0.00654 |
|  | OG0009672 | 0.02357 | 0.00186 | NA | 0.00037 | 0.00034 | 0.01165 | 0.02241 | NA | NA | 0.00493 | NA | NA |
|  | OG0009844 | 0.00478 | 0.00086 | 0.00379 | 0.00334 | 0.00269 | 0.00326 | 0.00151 | 0.00528 | 0.0019 | NA | 0.01051 | NA |
|  | OG0009968 | 0.00625 | 0.00125 | 0.018 | 0.0058 | 0.01427 | 0.00367 | NA | 0.01513 | 0.02135 | NA | 0.01017 | NA |
|  | OG0009970 | 0.04063 | 0.01511 | 0.01632 | 0.01735 | 0.01789 | 0.02648 | 0.00786 | 0.02972 | 0.0045 | NA | 0.04885 | 0.02556 |
|  | OG0010029 | 0.02609 | 0.00116 | NA | NA | 0.00155 | 0.01319 | NA | NA | 0.00501 | NA | NA | 0.02284 |
|  | OG0010055 | 0.02845 | NA | NA | 0.01049 | NA | 0.00264 | 0.00399 | NA | NA | NA | 0.0225 | NA |
|  | OG0010057_2 | NA | 0.00154 | NA | NA | NA | 0.00204 | NA | NA | 0.00392 | NA | NA | NA |
|  | OG0010088 | NA | 0.00228 | 0.02126 | 0.00345 | NA | 0.02313 | NA | NA | 0.02471 | 0.00844 | 0.01901 | NA |
|  | OG0010273 | NA | 0.00251 | 0.0012 | 0.00175 | 0.00366 | NA | 0.00221 | NA | 0.00664 | NA | NA | 0.00295 |
|  | OG0010322 | 0.00292 | 0.00402 | 0.01228 | 0.0043 | 0.00508 | 0.01063 | 0.00675 | 0.01011 | NA | 0.02967 | NA | NA |
|  | OG0010331 | 0.0247 | 0.00344 | 0.00343 | 0.00297 | 0.01802 | NA | NA | 0.00722 | NA | 0.01156 | 0.02311 | 0.00204 |
|  | OG0010665 | 0.0199 | 0.00391 | 0.00552 | 0.00578 | 0.00442 | 0.01279 | 0.0184 | 0.02017 | NA | NA | 0.00079 | NA |
|  | OG0010827 | 0.03869 | 0.02144 | 0.01424 | 0.02161 | 0.03132 | 0.01318 | 0.00104 | 0.02893 | 0.02135 | 0.01214 | 0.02609 | 0.04099 |
|  | OG0010927_1 | NA | 0.02525 | NA | 0.08119 | 0.00085 | NA | NA | NA | NA | NA | NA | NA |
|  | OG0010952 | NA | 0.03899 | 0.06765 | 0.03793 | 0.05284 | 0.04934 | 0.01695 | 0.0067 | 0.11239 | 0.1484 | 0.05419 | 0.01933 |
|  | OG0010985_1 | 0.00118 | 0.02284 | 0.02153 | 0.01565 | 0.02208 | 0.01243 | NA | 0.00151 | 0.04799 | 0.05263 | NA | 0.02577 |
|  | OG0011098 | NA | 0.00069 | 0.01847 | 0.00043 | 0.0143 | 0.0055 | 0.00869 | 0.0143 | 0.00167 | 0.01458 | NA | NA |
|  | OG0011168 | 0.01338 | 0.0004 | 0.00905 | 0.00407 | 0.00555 | 0.00595 | 0.00112 | 0.01139 | NA | NA | 0.00883 | 0.00628 |
|  | OG0011334 | 0.05498 | 0.00905 | 0.00957 | 0.00448 | 0.01809 | 0.0122 | 0.00667 | 0.01605 | 0.02254 | NA | 0.00455 | 0.01831 |
|  | OG0012114 | 0.00583 | 0.00536 | 0.02351 | 0.00785 | 0.00661 | 0.02059 | 0.01297 | 0.00638 | 0.02539 | 0.0109 | 0.01627 | NA |
|  | OG0012141 | NA | 0.02202 | 0.01948 | 0.05061 | 0.01065 | 0.00873 | NA | 0.01183 | 0.00885 | 0.05971 | 0.01421 | 0.03209 |
|  | average | 0.01910 | 0.00862 | 0.01668 | 0.01329 | 0.01277 | 0.01368 | 0.00816 | 0.01385 | 0.02049 | 0.03408 | 0.01933 | 0.01848 |
| *d_N_/d_S_* | OG0000093_1 | NA | 0.36701 | 0.1954 | 0.06463 | NA | 0.5315 | 2.0291 | NA | NA | NA | 0.0091 | 0.0029 |
|  | OG0000093_2 | 0.2789 | 0.20734 | 0.3048 | 0.0812 | 0.24498 | 0.312 | 0.0072 | 0.1672 | 0.1443 | 0.61998 | 0.3737 | 0.0015 |
|  | OG0008312 | 0.01404 | 0.09618 | 0.1088 | 0.05762 | 0.44107 | 0.03837 | 0.0906 | 0.0356 | 0.01746 | 0.14774 | 0.3788 | 0.12503 |
|  | OG0009404 | 0.3063 | 0.52633 | 0.13755 | 0.49828 | 0.25088 | 0.26283 | 0.40083 | 0.48967 | 0.0633 | 0.0767 | 0.4417 | 0.0806 |
|  | OG0009563 | 0.7038 | 0.0243 | 0.3497 | 0.34693 | 0.0714 | 0.0161 | NA | 0.00643 | NA | 0.1681 | NA | 0.4082 |
|  | OG0009672 | 1.20822 | 0.19653 | 0.1888 | 0.34128 | 1.39808 | 0.30175 | 0.16795 | 0.2843 | 0.0026 | 0.81145 | 0.3341 | 0.5162 |
|  | OG0009844 | 1.1972 | 0.27447 | 2.7592 | 0.35533 | 3.1146 | 1.45048 | 3.8507 | 0.5348 | 0.2547 | 0.4349 | 0.34043 | NA |
|  | OG0009968 | 2.3023 | 1.48559 | 0.14463 | 0.19964 | 0.26407 | 0.7511 | 0.0012 | 0.52002 | 0.55307 | 2.105 | 3.1203 | 1.2398 |
|  | OG0009970 | NA | 0.29556 | 0.14647 | 0.68167 | 1.17944 | NA | NA | 0.11828 | 0.0287 | 0.0071 | NA | 0.57413 |
|  | OG0009972 | 2.42418 | 1.02475 | 0.38936 | 0.23738 | 0.50338 | 0.61758 | NA | 1.5017 | 4.59943 | 0.55073 | 0.2218 | 0.3365 |
|  | OG0010029 | NA | 0.32587 | 0.08823 | 0.73088 | 0.86191 | 0.38045 | 0.06931 | 0.1985 | 0.04993 | 0.001 | 0.43338 | 0.32783 |
|  | OG0010055 | 0.10073 | 1.296 | 0.186 | 0.3195 | 0.73188 | 0.3983 | 1.34858 | 0.96549 | 0.008 | 0.4893 | 0.30643 | 2.03462 |
|  | OG0010057_1 | 0.4096 | 0.34472 | 0.46268 | 0.5918 | 1.98194 | 0.02777 | 0.2963 | 0.1993 | 0.3421 | 0.7295 | 0.0306 | 0.02582 |
|  | OG0010105 | 0.89377 | 1.65803 | 1.1404 | 0.295 | 0.8856 | 2.3535 | 1.936 | 1.15663 | 7.58964 | 0.5061 | 0.38432 | 1.6004 |
|  | OG0010246 | 8.5234 | 0.1993 | 13.9311 | 0.29533 | 0.67883 | 0.18087 | 4.67322 | 0.1087 | 0.0593 | NA | NA | 1.7445 |
|  | OG0010283 | 0.7252 | 0.51622 | 0.30565 | 0.2486 | 0.05475 | 0.55188 | 0.37663 | 0.1628 | 0.5238 | 0.3875 | 1.05097 | 0.001 |
|  | OG0010293 | 1.1985 | 0.29115 | 0.21594 | 0.6278 | 0.7902 | 0.54524 | 0.3836 | NA | 0.2286 | 1.6782 | 0.22678 | 0.35002 |
|  | OG0010891 | 5.044 | 0.5282 | 0.27977 | 8.9193 | 2.40544 | 0.93253 | 0.17274 | 0.4482 | 0.8511 | 1.2 | 0.00319 | 2.55617 |
|  | OG0010927_1 | NA | 0.9208 | 1.80322 | NA | NA | NA | NA | NA | NA | NA | NA | NA |
|  | OG0010952 | 20.68091 | 0.75868 | 0.28563 | 0.2748 | 0.97492 | 2.5135 | 0.00188 | 0.3705 | 1.4226 | NA | 0.3833 | 0.67335 |
|  | OG0010985_1 | 2.551 | 1.20066 | NA | 0.72414 | 2.05794 | 0.077 | 0.1141 | 0.38189 | NA | 1.26497 | 1.76475 | 1.6456 |
|  | OG0011080 | 0.42194 | 0.5303 | 1.9956 | 0.2303 | 0.1916 | 3.1132 | 0.0063 | 0.0354 | 0.18359 | NA | 1.2002 | 0.3485 |
|  | OG0011098 | 0.3233 | 0.53 | 1.2803 | 0.3433 | 2.44435 | 0.7233 | 0.28898 | 2.10263 | 1.29626 | 0.35629 | 0.0057 | 1.542 |
|  | OG0012141 | 0.4932 | 0.14296 | 0.5315 | 1.1334 | 0.33163 | 0.21972 | NA | 0.85973 | 0.76279 | 0.0217 | 0.07495 | 0.58463 |
|  | average | 2.49003 | 0.57254 | 1.18394 | 0.76513 | 0.99358 | 0.74086 | 0.85343 | 0.50704 | 0.94906 | 0.60822 | 0.55423 | 0.75997 |
